# Control of lipolysis by a population of oxytocinergic sympathetic neurons

**DOI:** 10.1101/2022.09.27.509745

**Authors:** Erwei Li, Luhong Wang, Daqing Wang, Jingyi Chi, Gordon I. Smith, Samuel Klein, Paul Cohen, Evan D. Rosen

## Abstract

Oxytocin (OXT), a nine amino acid peptide produced in the hypothalamus and released by the posterior pituitary, has well-known actions in parturition, lactation, and social behavior^1^, and has become an intriguing therapeutic target for diseases like autism and schizophrenia^2^. Exogenous OXT has also been shown to promote weight loss, among other beneficial metabolic effects^1,3^, suggesting that its therapeutic potential may extend to diabetes and obesity^1,4^. It is unclear, however, whether endogenous OXT participates in metabolic homeostasis. Here we show that OXT is a critical regulator of adipose tissue lipolysis in both mice and humans. In addition, OXT serves to license the ability of β- adrenergic agonists to fully promote lipolysis. Most surprisingly, the relevant source of OXT in these metabolic actions is a previously unidentified subpopulation of tyrosine hydroxylase (TH)-positive sympathetic neurons. Our data reveal that OXT from the peripheral nervous system is an endogenous regulator of adipose and systemic metabolism.

---

The ability of exogenous OXT to promote weight loss in rodents has primarily been attributed to actions on food intake, but recent papers have called this idea into question^5-7^. Other studies have suggested that OXT has effects on peripheral metabolism, including some dating back to the early 1960’s showing that OXT administration increases serum free fatty acids (FFAs) in peripartum women^8^. Oxytocin receptor (OXTR) is found in high abundance on the surface of adipocytes, and global OXTR knockout mice show mild, late onset obesity^9^, suggesting that OXT is an endogenous regulator of metabolism, although the extent of its contribution is still unclear.

## OXT is a direct inducer of lipolysis in adipocytes

To investigate the direct actions of OXT on adipose tissue, we first assessed OXTR expression in three different murine adipose depots. White adipose tissue (WAT), especially visceral, has the highest *Oxtr* mRNA and protein levels (**Extended Data Fig. 1a, b**), with expression highest in the adipocytes themselves (**Extended Data Fig. 1c, d**). Consistent with this, *Oxtr* expression increased during adipose differentiation (**Extended Data Fig. 1e, f**). Interestingly, cold exposure caused *Oxtr* expression to rise in whole epididymal (eWAT) and inguinal (iWAT) adipose tissue and in isolated adipocytes (**Extended Data Fig. 1g, h**).

To confirm direct action of OXT on adipocytes, we differentiated mouse adipocytes from SVF and then treated with OXT, which increased glycerol release by 1.6-fold (**Fig. 1a, Extended Data Figure 2a**), with a similar effect seen in human adipose explants (**Fig. 1b**), confirming direct, though weak, lipolytic action of OXT. We also tested the effect of OXT on lipolysis *in vivo*; administration of OXT to WT mice increased serum glycerol and free fatty acids (FFA) to a degree comparable to that seen *in vitro* (**Fig. 1c**).

**Figure 1.**
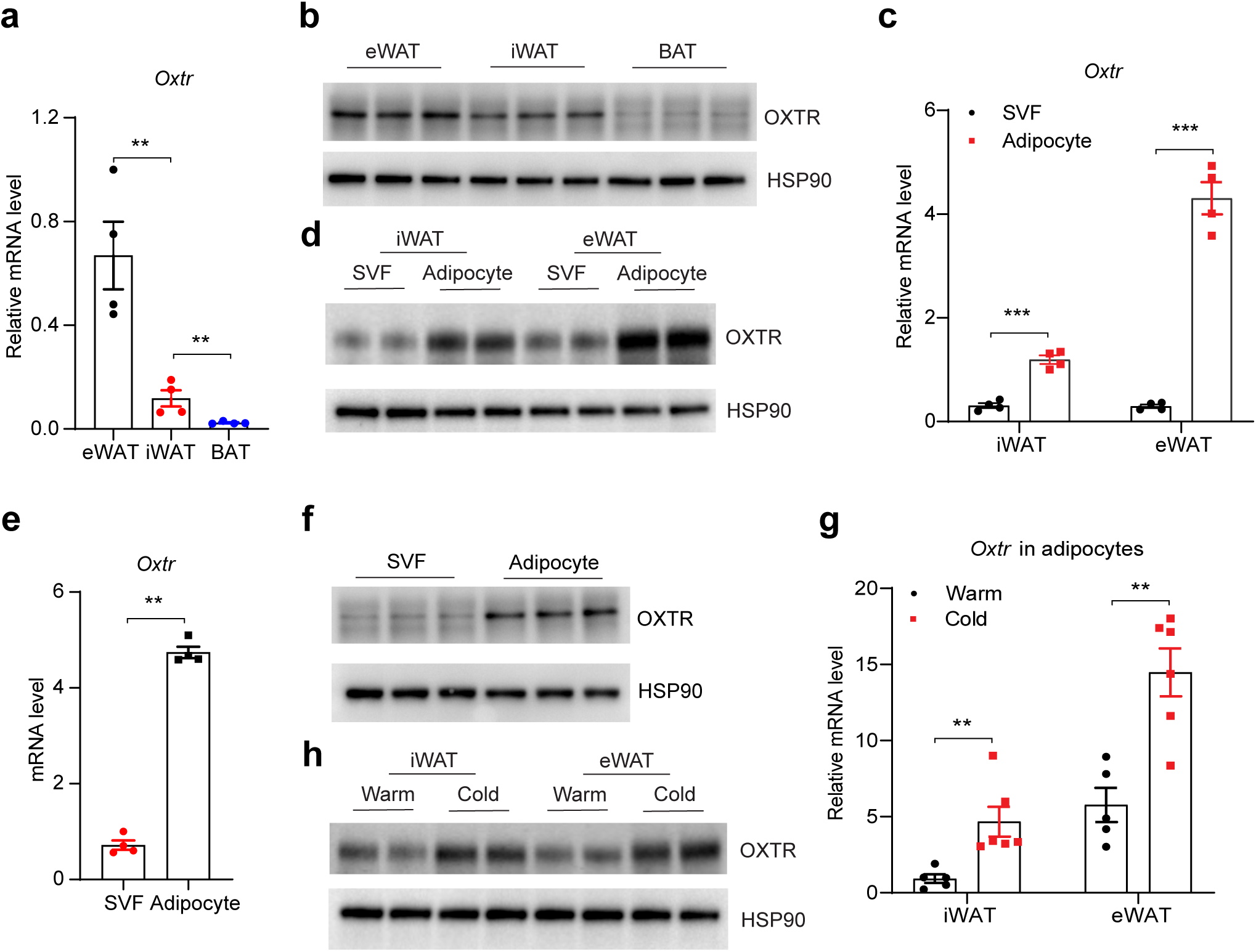
OXT promotes lipolysis in vitro and in vivo and licenses beta-adrenergic lipolysis. **a.** Glycerol release from mouse adipocytes differentiated from SVF and treated with OXT (10uM) for 3 hours; n=4. **b.** Glycerol release from human adipose explants treated with OXT (10uM) overnight; n=4. **c.** Serum glycerol and FFA levels in WT mice before and 1 hour after intraperitoneally OXT injection (3 mg/kg body weight); n=5. **d.** Serum glycerol and FFA levels from *Oxtr*^flox^ and *Oxtr*^ΔAd^ mice fasted overnight or fed *ad libitum*; *Oxtr*^flox^ n=9, *Oxtr*^ΔAd^ n=8. **e.** Serum glycerol and FFA levels from *Oxtr*^flox^ and *Oxtr*^ΔAd^ mice housed at 30°C vs. 4°C overnight; n=6. **f.** Glycerol release from cultured adipocytes treated with OXT (10uM), isoproterenol (ISO; 10uM), or both OXT and ISO, in the presence or absence of the MEK inhibitor Trametinib (Tra)(5 nM) or the ERK inhibitor Temuterkib (Tem)(2 uM); n=4. **g.** Glycerol release from cultured adipocytes treated with varying doses of ISO in the presence or absence of OXT (10uM); n=4. **h.** Glycerol and FFA levels of WT mice 1 hour after IP injection of OXT (3mg/kg body weight), ISO (10mg/kg body weight), or both; n=5. **i.** Glycerol and FFA levels of WT mice before and 1 hour after the IP injection of ISO (10mg/kg body weight) with and without atosiban (5mg/kg body weight); n=5. **j.** Serum glycerol and FFA levels after ISO treatment of *Oxtr*^flox^ and *Oxtr*^ΔAd^ mice; n=6. Data are presented as mean ± s.e.m. Statistical comparisons were made using 2-tailed Student’s t test (a-f, h, i) or 2-way ANOVA (g, j). *P < 0.05; **P < 0.01.

The classical signaling cascade leading to lipolysis involves engagement of β-adrenergic receptors by norepinephrine, followed by Gsα activation, cAMP accumulation, and induction of protein kinase A (PKA) activity. PKA then phosphorylates hormone-sensitive lipase (HSL) and perilipin 1 (PLIN1)^10^. An alternate, but related, pathway involves activation of protein kinase G (PKG) by natriuretic peptides, at least in human WAT^11^. OXT signaling generally involves activation of Gq and not Gsα^12^, and would therefore not be expected to utilize this classical pathway. Consistent with this, we did not observe OXT-dependent activation of PKA or cAMP accumulation in adipocytes (**Extended Data Fig. 2b, c**). Similarly, inhibition of PKA and (to a lesser extent) PKG blocked the ability of the β-adrenergic agonist isoproterenol (ISO) to promote lipolysis, but had no effect on OXT-induced lipolysis (**Extended Data Fig. 2d, e**). In some tissues, OXT has been shown to signal via the MAPK-ERK pathway^12,13^, and ERK has been proposed to regulate HSL activity and lipolysis^14,15^. We therefore assessed whether OXT activates ERK in adipocytes and found that it does so in a time- and dose-dependent manner (**Extended Data Fig. 2f, g**). Consistent with the notion of an ERK-dependent lipolytic pathway downstream of OXTR, the MEK inhibitor Trametinib (Tra) and the ERK inhibitor Temuterkib (Tem) both reduce OXT-induced glycerol release (**Extended Data Fig. 2h**).

To further investigate the physiological role of OXT signaling on lipolysis *in vivo,* we crossed *Oxtr*^flox^ mice^16^ and Adipoq-Cre mice^17^ to generate adipocyte-specific *Oxtr* knockout mice (hereafter referred to as *Oxtr*^ΔAd^) (**Extended Data Fig. 3a-c**). These mice have no alteration in body weight or body composition on chow diet, and gain weight normally after high fat feeding (**Extended Data Fig. 3d, e**). Although there is no difference in total body weight after 16 weeks of HFD, *Oxtr*^ΔAd^ mice have increased eWAT mass and reduced liver weight, consistent with reduced lipolysis in eWAT ultimately reducing hepatic lipid content (**Extended Data Fig. 3f, g).**

Fasting provoked reduced levels of FFA and glycerol in the serum of chow-fed *Oxtr*^ΔAd^ mice relative to control mice (**Fig. 1d**). Similarly, cold exposure, another lipolytic stimulus, led to a similar discrepancy between *Oxtr*^ΔAd^ and control mice (**Fig. 1e**). Consistent with this, *Oxtr*^ΔAd^ mice were unable to defend their body temperature in response to a cold challenge despite moderately elevated UCP-1 protein levels in their iWAT, reflecting the reduced fatty acid substrate required to fuel thermogenesis under these conditions (**Extended Data Fig. 3h, i**). No direct effect of OXT administration on thermogenic gene expression was noted in adipocytes. (**Extended Data Fig. 3j, k**).

## OXT licenses the actions of **β**-agonist on lipolysis

Cold exposed *Oxtr*^ΔAd^ mice have normal levels of total hormone-sensitive lipase (HSL) and adipose triglyceride lipase (ATGL) protein in eWAT and iWAT, but display significantly reduced pHSL and pPLIN1 levels at PKA sites after cold exposure (**Extended Data Fig. 4a, b**). These data suggest that loss of OXT-OXTR signaling compromises the ability of catecholamines, which are secreted by sympathetic neurons directly into the fat pad, to activate the lipolytic machinery. This prompted us to explore the relationship between OXT and β-adrenergic agents with respect to lipolysis. We noted a synergistic effect between OXT and ISO in promoting lipolysis in mouse adipocytes, an effect seen across a broad range of ISO concentrations (**Fig. 1f, g**). Dose-response experiments demonstrated that OXT not only increases catecholamine sensitivity but also the maximal capacity of ISO to induce lipolysis (**Fig. 1g**). MEK/ERK inhibition minimally weakened the effect of ISO by itself, but completely abrogated the ability of OXT to enhance ISO-mediated lipolysis (**Fig. 1f**). Importantly, these results were recapitulated in human adipose explants (**Extended Data Fig. 4c, d**).

We next sought additional evidence that OXT enables or “licenses” the full lipolytic effect of β-agonists *in vivo*. First, we tested whether exogenous OXT could enhance the actions of co-administered ISO, and found that, as before, OXT is itself a relatively weak lipolytic inducer, but a potent enhancer of catecholamine-induced lipolysis (**Fig. 1h**). Conversely, the OXTR antagonist Atosiban significantly reduced ISO-induced lipolysis in mice (**Fig. 1i**). Similarly, *Oxtr*^ΔAd^ mice display a blunted response to ISO (**Fig. 1j**). These data indicate that OXT-OXTR signaling is both necessary and sufficient to license the full potential of β-adrenergic agonists to exert their full effect on lipolysis. They also suggest that OXT is an endogenous regulator of lipolysis.

At a molecular level, OXT increased the ability of ISO to promote PKA-mediated phosphorylation of PLIN1 (but not HSL), and this effect was blocked by MEK/ERK inhibition (**Extended Data Fig. 4e**). OXT enhances PKA-mediated PLIN1 phosphorylation without increasing cAMP levels or the ability of PKA to phosphorylate its substrate proteins or peptides (**Extended Data Fig. 4e, f**), suggesting that OXT and MAP/ERK signaling modify specific targets, making them more amenable to binding or phosphorylation by PKA. This notion is consistent with previously described ERK-mediated phosphorylation of HSL^18^.

One effect of PKA-mediated phosphorylation of HSL is to induce translocation to the lipid droplet^10,19^. We found that this process is unaffected by OXT, either alone or in combination with ISO (**Extended Data Fig. 5a**). Similarly, ISO reduces total PLIN1 levels and alters its distribution on the surface of the lipid droplet, which is believed to help expose lipid to the actions of HSL^20-22^. We confirmed these actions of ISO, but OXT, either alone or in combination with ISO, had no additional effect (**Extended Data Fig. 5b, c**). In addition, we found that ATGL is likely not involved in the effect of OXT on ISO-induced lipolysis, as the ATGL inhibitor Atglistatin reduces overall lipolysis but does not block the ability of OXT to enhance the effect of β-agonist. (**Extended Data Fig. 5d, e**).

## Peripheral sympathetic neurons are the relevant source of OXT in WAT

Given that OXT signaling is required for lipolysis associated with fasting and cold exposure, we speculated that OXT levels in serum would increase under these conditions, but this was not what we observed (**Fig. 2a, b**). Surprisingly, however, intra-adipose tissue levels of OXT were significantly elevated after both cold exposure and fasting (**Fig. 2a, b**). This phenomenon is not restricted to mice, as humans subjected to an overnight fast exhibit elevated serum OXT levels in WAT, but not blood, compared to three hours post-refeeding (**Fig. 2c, d**). This decoupling of serum and tissue OXT levels suggested to us that there might be a unique source of OXT in fat. Assessment of our single nucleus RNA-seq data of from mouse and human adipose tissue^23^, however, revealed few if any OXT-expressing cells within murine or human WAT (**Extended Data Fig. 6a, b**). Concerned that this might reflect low levels of OXT expression that might be hard to detect using single cell strategies, we crossed Oxt-Cre mice to Ai9 mice, which express tdTomato in a Cre-dependent manner. Adipose SVF harvested from Oxt-Cre::Ai9 mice were sorted by FACS, but no tdTomato-positive cell population was identified in either iWAT or eWAT (**Extended Data Fig. 6c**). Importantly, neurons that innervate the fat pad are not detectable with single cell or FACS-based approaches, as their cell bodies reside in ganglia located outside of the adipose depot. Given that fasting and cold both increase sympathetic activity in WAT^24^, we considered that sympathetic nerves might be the source of intra-adipose OXT. Of note, expression of OXT outside of the CNS has been reported in some sensory neurons and in the intrinsic nerves of the gut^25,26^, but has not been reported in the sympathetic nervous system (SNS). To assess this possibility, we crossed Oxt-Cre mice to a reporter line expressing GFP and mCherry in a Cre-dependent manner (H2B-TRAP)^27^, and visualized the location of GFP using Adipo-Clear^28^. GFP was found to co-localize with a subset of tyrosine hydroxylase (TH)-positive nerves (**Fig. 2e**). Sympathetic ganglia that innervate WAT in mice include paravertebral chain ganglia at T12-L2 (for iWAT)^29^ and the aorticorenal ganglion (ARG; for eWAT)^30^. In Oxt-Cre::Ai9 mice, we consistently visualized tdTomato-positive sympathetic neurons in the ARG (**Fig. 2f**). Immunostaining of the ARG from these mice demonstrated co- localization of TH and tdTomato (**Fig. 2g**). Similarly, Oxt-Cre::H2B-TRAP mice showed GFP and mCherry expression in L1 chain ganglion, but not in ganglia that do not innervate adipose tissue, such as at T5; these cells also exhibit co-localization of GFP and TH staining (**Extended Data Fig. 7a, b**). This indicates that *Oxt* expression (driving Cre) had occurred at some point in the developmental history of the sympathetic neurons in ganglia that innervate WAT, but it does not prove that expression is being driven from the *Oxt* locus in the adult mouse. Furthermore, because these ganglia innervate many other tissues and organs, it remained possible that OXT expression was occurring specifically in sympathetic neurons that do not innervate WAT. To address these issues more definitively, we injected a retrograde adeno-associated virus (AAV) expressing Cre-inducible mCherry directly into the eWAT of Oxt-Cre mice. The resulting mCherry+ TH+ neurons in the ARG (**Fig. 2h**) indicate that sympathetic neurons innervating eWAT actively express OXT in the adult animal. Using the same approach, we found that sympathetic neurons innervating iWAT (e.g., in L1 ganglia) also actively express OXT (**Extended Data Fig. 7c, d**).

**Figure 2.**
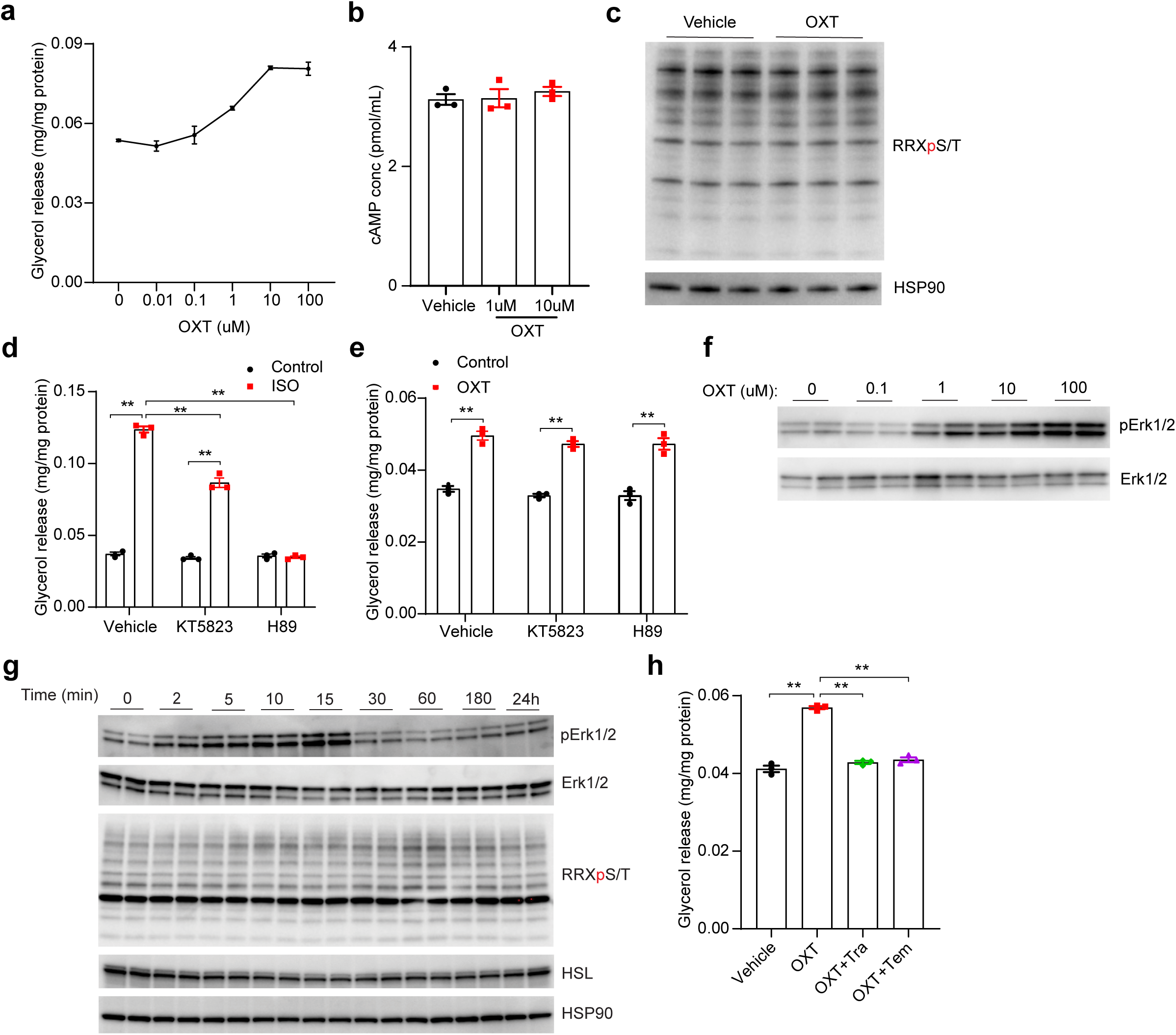
Sympathetic neurons are the source of OXT within adipose tissue. **a.** Serum and intra-adipose tissue levels of OXT in eWAT and iWAT of WT mice housed at 30°C vs. 4°C for 1 week; Serum OXT n=6-8, WAT OXT n=5. **b.** Serum and intra-adipose tissue levels of OXT in eWAT of WT mice after feeding vs. fasted overnight; n=5. **c.** Schematic of serum and adipose tissue sample collection from human patients. **d.** Serum and intra-adipose tissue levels of OXT in human patients fasted overnight vs 3 hours after refeeding; n=10. **e.** Sympathetic nerves from cleared eWAT of Oxt-Cre::H2B-TRAP mice. GFP (green) marks expression of Oxt-Cre, with counterstaining with anti-TH (red). Scale bar, 100 um. **f.** Aorticorenal ganglion (ARG) of *Oxt*-Cre::Ai9 mice perfused with PBS. tdTomato (red) marks expression of Oxt-Cre. Arrowhead indicates autofluorescent blood vessel debris. Scale bar, 200 um. **g.** Co-immunostaining of tdTomato and TH in ARG of *Oxt*-Cre::Ai9 mice. Scale bars, 50 um. **h.** Co-immunostaining of mCherry and TH in ARG of *Oxt*-Cre mice injected with AAV2/retro- syn-Flex-mCherry in eWAT. Scale bars, 50 um. Data are presented as mean ± s.e.m. Statistical comparisons were made using 2-tailed Student’s t test (a, b) or 2-tailed paired Student’s t test (d). *P < 0.05; **P < 0.01.

## Activation of the SNS promotes OXT release

We next asked whether activation of the sympathetic nervous system (SNS) is sufficient to cause OXT release from neurons innervating WAT. Stereotaxic injections of AAV8-hSyn-DIO- hM3Dq-mCherry were made into the raphe pallidus (RPa) of 6-8-week-old male Vglut3-IRES- Cre mice, thus expressing the DREADD hM3Dq in Vglut3+ neurons, which are afferents of the sympathetic response^31^ (**Fig. 3a and Extended Data Fig. 7e**). Clozapine-N-oxide (CNO) or saline were delivered using minipumps implanted subcutaneously. CNO caused increased serum glycerol levels in mice with DREADD expression, indicating that raphe activation led, as expected, to higher SNS activity with subsequent lipolysis (**Fig. 3b**). Notably, raphe activation resulted in increased OXT levels in iWAT and eWAT, but not in serum (**Fig. 3c-e**). In order to assess the effect of stimulating sympathetic nerve terminals in WAT directly, we crossed Oxt- Cre mice to Ai32 mice, which express channelrhodopsin in a Cre-dependent manner. eWAT explants were harvested from these mice and pulsed with blue light, which induced a significant release of OXT into the medium (**Fig. 3f**), further corroborating the notion that direct activation of sympathetic nerve terminals within adipose tissue leads to local OXT release.

**Figure 3.**
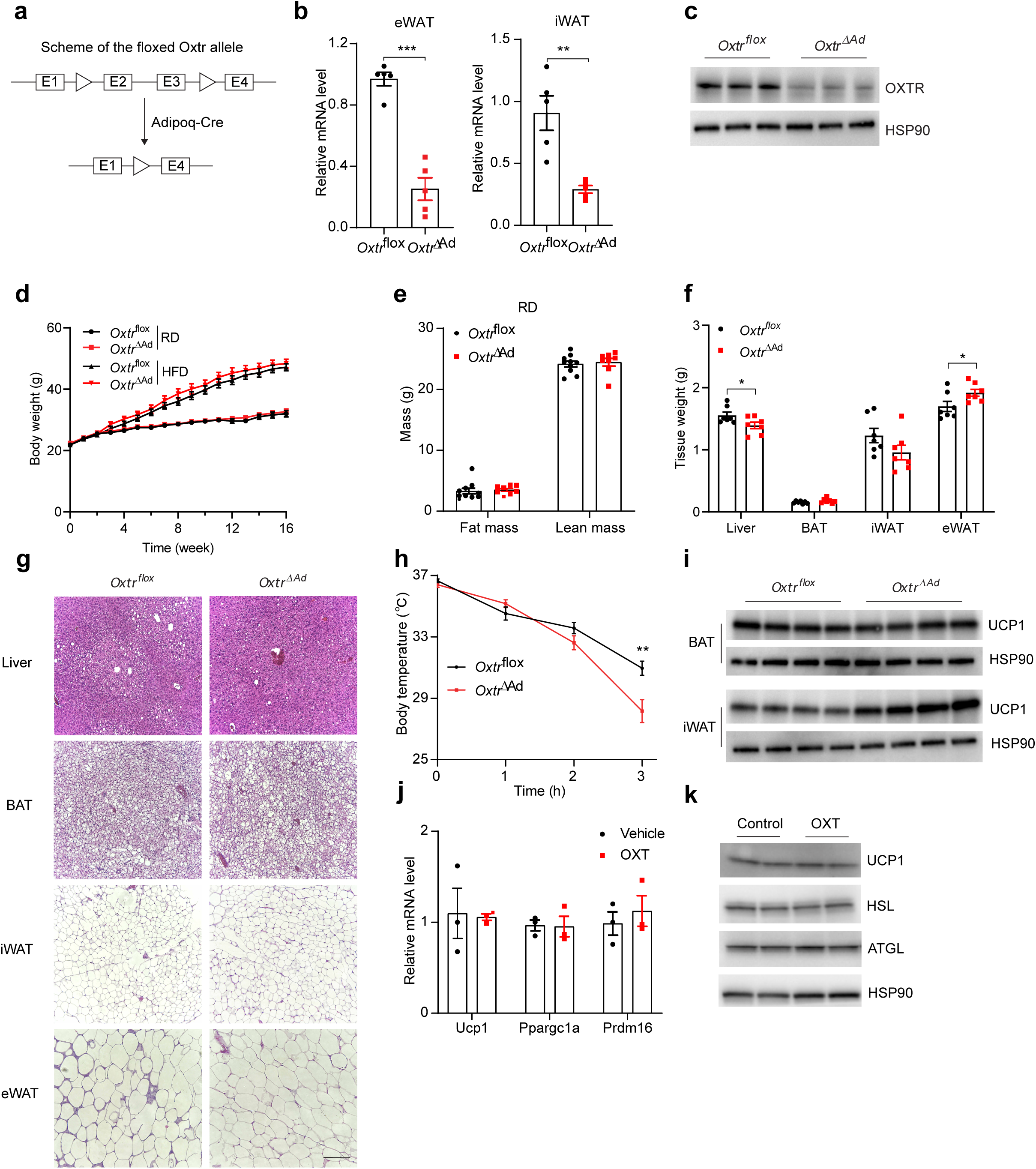
SNS activation increases tissue OXT levels. **a.** Schematic of experiment testing effect of chemogenetic activation of glutamatergic neurons in the raphe pallidus (RPa) on OXT levels in serum and WAT. RPa of Vglut3-IRES-Cre mice were injected with AAV2/8-hSyn-DIO-hM3Dq-mCherry virus (or AAV2/8-hSyn-DIO-mCherry as control), which expresses the DREADD hM3Dq in a Cre-dependent manner. Administration of CNO activates the DREADD, promoting activation of the SNS from the RPa, through the intermediolateral nucleus of the spinal cord (IML). **b.** Serum glycerol levels in mice from a treated with CNO or vehicle; n=4-5. **c.** Serum OXT levels in mice from a treated with CNO or vehicle; n=4-5. **d.** Tissue OXT levels in iWAT of mice from a treated with CNO or vehicle; n=4-5. **e.** Tissue OXT levels in eWAT of mice from a treated with CNO or vehicle; n=4-5. **f.** Schematic of experiment testing effect of optogenetic activation of peripheral sympathetic nerves on OXT release. eWAT from Oxt-Cre::Ai32 mice expressing channelrhodospin in a Cre- dependent manner was harvested, cultured in medium, and exposed to blue light pulses ex vivo. OXT protein levels in medium was determined by ELISA; n=5. Data are presented as mean ± s.e.m. Statistical comparisons were made using 2-tailed Student’s t test (b-f). *P < 0.05; **P < 0.01.

## OXT from TH-positive sympathetic neurons promotes lipolysis

To assess whether OXT released from sympathetic neurons plays a functional role in adipose tissue, we generated *Oxt*^flox^ mice (**Extended Data Fig. 8a**) and crossed them to Th-Cre mice, generating animals that lack OXT in sympathetic neurons (*Oxt*^ΔTH^). These mice are viable and display no difference in food intake or body weight on chow or high fat diet (**Extended Data Fig. 8b-c**). As predicted, these mice exhibit reduced OXT levels in WAT, but not in the serum (**Fig. 4a, b**). These mice also display reduced lipolysis in response to exogenous ISO, fasting and cold exposure (**Fig. 4c-e**). Consistent with this, they were also unable to defend their body temperature when placed in the cold (**Fig. 4f**). This is highly unlikely to be due to inadvertent knockout of OXT in the hypothalamus, because (a) immunostaining of TH and OXT in hypothalamus reveals minimal overlap at the protein level (**Extended data Fig. 8d**); (b) serum OXT levels are unaffected in *Oxt*^ΔTH^ mice (**Fig. 4a**); and (c) immunostaining for OXT in the hypothalamus of *Oxt*^ΔTH^ mice does not suggest reduced Oxt expression (**Extended data Fig. 8e**). However, it remained possible that the effect on WAT lipolysis could be due to loss of *Oxt* expression in TH-positive sympathetic neurons that innervate other organs, which might then affect WAT indirectly. To prove that this is not the case, we injected a retrograde AAV2/Retro- hSyn-Cre virus (or AAV2/Retro-hSyn-mCherry control virus) directly into the inguinal fat pads of *Oxt*^flox^ mice. These viruses travel up the sympathetic axons that innervate the iWAT, but do not cross the synapse. This manipulation is designed to delete the *Oxt* gene specifically in the sympathetic neurons innervating the fat pad receiving the AAV-retro-Cre virus. Three weeks after injection, fat pads were excised and treated as explants with isoproterenol. Adipose tissue innervated by sympathetic neurons that received the Cre expressing virus were deficient in their ability to release glycerol in response to β-agonist (**Fig. 4g**). Taken together, our data support a significant lipolytic role for a subpopulation of oxytocinergic TH+ sympathetic neurons.

**Figure 4.**
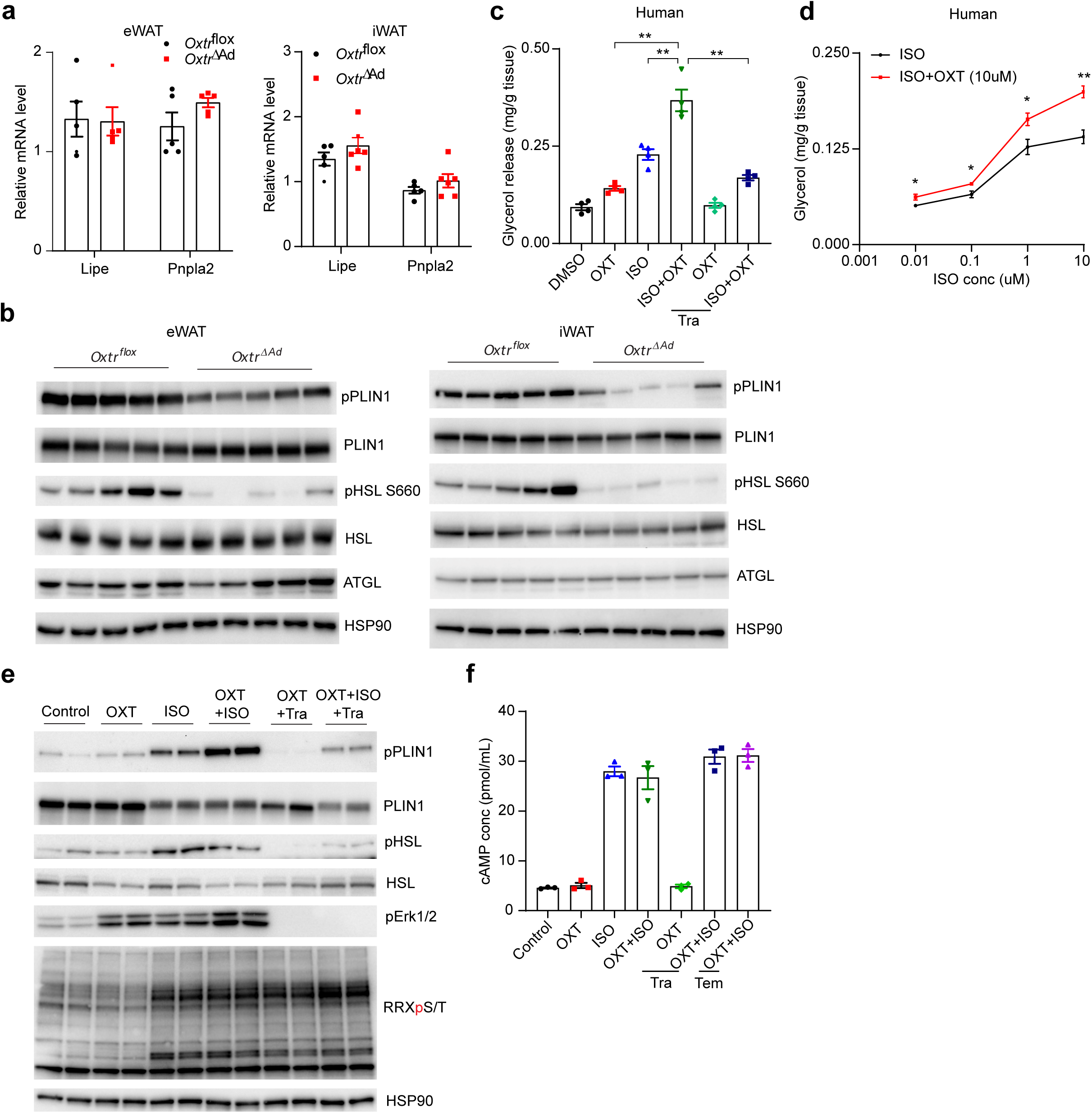
Loss of OXT in TH+ neurons reduces lipolysis. **a.** Serum OXT levels of cold-challenged (24 hours) *Oxt*^flox^ and *Oxt*^ΔTH^ mice; *Oxt*^flox^ n=5, *Oxt*^ΔTH^ n=6. **b.** Adipose tissue (eWAT) OXT levels of cold-challenged (24 hours) *Oxt*^flox^ and *Oxt*^ΔTH^ mice; *Oxt*^flox^ n=5, *Oxt*^ΔTH^ n=6. **c.** Glycerol and FFA levels in *Oxt*^flox^ and *Oxt*^ΔTH^ mice following cold exposure (24 hours); n=6. **d.** Glycerol and FFA levels in *Oxt*^flox^ and *Oxt*^ΔTH^ mice following overnight fasting; *Oxt*^flox^ n=8, *Oxt*^ΔTH^ n=7. **e.** Serum glycerol and FFA levels after ISO treatment of *Oxt*^flox^ and *Oxt*^ΔTH^ mice; n=5. **f.** Rectal temperature of *Oxt*^flox^ and *Oxt*^ΔTH^ mice exposed to 4°C; n=7. **g.** Glycerol release from the cultured iWAT explants of *Oxt*^flox^ mice injected with AAV2/Retro- hSyn-Cre or AAV2/Retro-hSyn-mCherry after ISO treatment; n=5. Data are presented as mean ± s.e.m. Statistical comparisons were made using 2-tailed Student’s t test (a-d) or 2-way ANOVA (e-g). *P < 0.05; **P < 0.01.

## Discussion

Adipose tissue triglycerides are the body’s major fuel reserve, and the release of FFA from adipose tissue is critical for survival during periods of food deprivation and for muscle function during endurance exercise^32,33^ However, an excessive rate of lipolysis and increased plasma FFA concentrations can have adverse effects on metabolic health by causing insulin resistance in liver and skeletal muscle and by altering β-cell glucose stimulated insulin secretion^34-36^. Despite being one of the most well-studied metabolic pathways, new insights into the genetics, biochemistry, cell biology, and physiology of lipolysis continue to accumulate^10^. The canonical pathway of homeostatic lipolysis involves release of norepinephrine from TH-positive sympathetic neurons that innervate the fat pad; norepinephrine activates β-adrenergic receptors on the surface of the adipocyte, triggering an intracellular signaling cascade that involves cAMP accumulation, PKA activation, and phosphorylation of key actors like HSL and PLIN1. Our data indicate that this picture is incomplete in that the ability of β-adrenergic agonists to maximally stimulate lipolysis requires the presence of an active OXT-OXTR-ERK signaling cascade.

Perhaps the most surprising result of our work is the identification of a subpopulation of oxytocinergic TH+ sympathetic neurons that innervate WAT and account for the observed metabolic effects on adipocytes. OXT expression has been reported in some sensory nerves and in the enteric nervous system, but not in the SNS, nor has its functional role in these peripheral sites been delineated.

Limitations of our study include the fact that we do not exclude a role for hypothalamic OXT on lipolysis. Further studies using our newly developed *Oxt*^flox^ mice may shed light on this issue. Furthermore, although we have identified ERK-mediated enhancement of PKA activity on specific substrates (e.g., HSL and PLIN1) as a pathway by which OXT enhances lipolysis, we have not defined a precise mechanism by which this occurs. Finally, our data were primarily collected in mice, but we note that the ability of OXT to increase lipolysis in human adipocytes and the elevation of OXT in WAT, but not serum, of fasted subjects suggests that this system is also operative in people.

Collectively, we identify a role for the local elaboration of OXT within white adipose depots to regulate lipolysis. These sources and actions of OXT will be important to keep in mind as the next generation of OXT-based therapeutics are developed for obstetric, psychiatric, and metabolic use^2,37^.

## METHODS

### Animals

Animal care and experiments were performed with approval from the BIDMC Institutional Animal Care and Use Committee. Mice were housed on a 12-h light/dark cycle at constant temperature (23°C) with free access to food and water. C57B6/J mice were purchased from Jackson Laboratory and used for experiments after an acclimatization period of several weeks. Experiments involving adipocyte-specific *Oxtr* knockout mice and sympathetic neuron-specific *Oxt* mice used littermate controls. For studies in lean mice, 8–12-week old male mice were used. The chow diet used was purchased from Harlan Teklad (catalog *8664; 12.5 kcal from fat); high fat diet was from Research Diets (catalog* D12492i; 60 kcal from fat). For DIO mice, chow diet was replaced with high fat diet beginning at 6 weeks of age. To label OXT expressing cells, H2B-TRAP mice27 (Jax 029789) or Ai9(RCL-tdT) mice (Jax 007909) were crossed with Oxt-Cre mice (Jax 024234). Other mouse strains used include: Vglut3-IRES-Cre (Zeng et al., 2019) and Ai32 mice (Jax 024109). Oxtrflox mice (Jax 008471) were crossed to Adipoq-Cre mice (Jax 028020) to generate adipocyte-specific Oxtr mice. Oxtflox mice were generated using CRISPR-Cas9. Two guide RNAs (GGGCCTGCCTCTAAACAGCG, GCTCCCTCTTGACGCCGTGA) flanking the first exon of Oxt were synthesized by PNA-BIO. The CAS9 enzyme (PNA-BIO, CP01), the guide RNAs and the single-stranded DNA repair template (synthesized by Genewiz) were microinjected together into fertilized eggs of FVB/NJ mice at the transgenic core of Beth Israel Deaconess Medical Center. F1 progeny were genotyped with the following primers: GATGACCTTGACCCTAGCCC, CGAGGTCAGAGCCAGTAAGC. The wild-type allele yielded a band of 263 bp and the Nts-floxed allele yielded a band of 297 bp. F1 progeny carrying the *Oxt*^flox^ allele were then backcrossed to C57BL/6J mice purchased from Jackson Laboratory for 10 generations to create congenic mice. Progeny carrying the *Oxt*^flox^ allele were then crossed to Th-Cre mice (European Mouse Mutant Archive; EM:00254) to generate sympathetic neuron-specific *Oxt* knockout mice.

### Effect of OXT on lipolysis in human samples

Abdominal and perianal subcutaneous adipose tissue was used to test the effect of OXT, ISO and ISO+OXT, with or without Trametinib, on lipolysis. Human adipose tissue was collected under Beth Israel Deaconess Medical Center Committee on Clinical Investigations IRB 2011P000079. Written informed consent was obtained from each individual (n = 4) donating tissue and samples were anonymized and handled according to the ethical guidelines set forth by the BIDMC Committee on Clinical Investigations. Subjects were recruited from the plastic surgeon operating room schedule at BIDMC in consecutive fashion, as scheduling permitted. Male and female subjects over the age of 18 undergoing elective plastic surgery procedures and free of other acute medical conditions were included. Subjects taking insulin-sensitizing medications such as thiazolidinediones or metformin, chromatin-modifying drugs such as valproic acid, and drugs known to induce insulin resistance such as mTOR inhibitors (for example, Sirolimus or Tacrolimus) or systemic steroids were excluded. The samples were verified as tumor free by gross pathological assessment. BMI measures were derived from electronic medical records and confirmed by self-reporting.

### Effect of feeding on plasma and adipose tissue OXT in humans

The effect of mixed meal ingestion on plasma OXT concentrations and adipose tissue OXT content were evaluated in 9 women and 1 man (mean age: 44 ± 3 years old; body mass index: 44 ± 2 kg/m^2^). Written, informed consent was obtained from all subjects before their participation in this study, which was approved by the Human Research Protection Office at Washington University School of Medicine in St. Louis, MO and registered in ClinicalTrials.gov (NCT03091725). All participants completed a screening evaluation that included a medical history and physical examination, and standard blood tests. Potential participants who had a history of diabetes or other serious chronic diseases, were taking medications that could affect study outcome measures, or consumed excessive amounts of alcohol (>21 drinks per week for men and >14 drinks per week for women) were excluded. Approximately 1 week later, subjects were admitted to the Washington University School of Medicine Clinical Translational Research Unit in the afternoon and consumed a standard evening meal. The following morning, after participants fasted for ∼11 hours overnight, a blood sample was obtained to assess fasting plasma OXT concentration and subcutaneous abdominal adipose tissue was obtained from the periumbilical area by aspiration through a 3-mm liposuction cannula (Tulip Medical Products, San Diego, CA) to assess fasting adipose tissue OXT content. At 0830 h, subjects ingested a liquid meal (containing 50 g glucose, 18 g fat, and 22 g protein), which was provided in seven equal aliquots every 5 minutes over 30 minutes. Blood and adipose tissue samples were obtained again at 3 hours after initiating meal ingestion to assess postprandial plasma OXT concentration and adipose tissue OXT content.

### Oxytocin measurement

For serum OXT measurements, mouse whole blood was collected into a capillary blood collection tube (Thermo Fisher Scientific, NC9141704) and spun at 1500g for 15LJmin to remove cells. Supernatant was aliquoted and stored at −80LJ°C. Mouse and human serum OXT concentration was measured using an ELISA kit (Enzo Life Sciences, ADI-900-153A-0001) according to the manufacturer’s instructions. For measurement of oxytocin in adipose tissue of mice and humans, 50mg adipose tissue was homogenized in PBS containing protease inhibitor cocktail (Promega, G6521) in a Qiagen TissueLyser II (85300). Lysates were centrifuged at 12,000g for 15 minutes and subnatant was collected using 500ul syringes (Thermo Fisher Scientific, 14-826-79). Oxytocin levels in the subnatant were measured using an ELISA kit (Enzo Life Sciences, ADI-900-153A-0001) according to the manufacturer’s instuctions; results were normalized to tissue weight.

### Mouse SVF isolation and adipocyte fraction

Inguinal adipose tissue dissected from 8-12-week-old C57BL/6J male mice, minced and digested in PBS with 10mM CaCl2, 1.5U/ml collagenase D (Sigma, 11088882001) and 2.4U/ml dispase II for 30 min at 37°C. The cell suspension was then filtered through a 100um cell strainer, centrifuged at 800 rpm for 5 min. Floating adipocytes were collected for RNA or protein extraction, while the cell pellet was lysed by ACK buffer (Thermo Fisher), resuspended in DMEM medium with 10% FBS and 1% P/S and centrifuged at 800 rpm for 5 min again. This pellet was lysed with Trizol for RNA isolation or with RIPA buffer for immunoblotting, or resuspended and plated prior to differentiation.

### Primary adipocyte culture

Adipocyte differentiation was triggered by treating confluent SVF cells with induction DMEM medium containing 10% FBS, 1% P/S, 0.5 mM isobutylmethylxanthine, 125 nM indomethacin, 2 µg/ml dexamethasone, 850 nM insulin, 1 nM T3 and 0.5 µM rosiglitazone (all from Sigma). Two days later, cells were switched to maintenance medium containing 10% FBS, 1% P/S, 850 nM insulin and 1 nM T3. In some experiments, primary adipocytes were transduced with adenoviral constructs (Cre-GFP Adenovirus, #000023A; GFP Adenovirus, #000541A, both from Applied Biological Materials).

### Metabolic studies

Six-week-old male mice were fed a standard diet (chow) or high-fat diet (Research Diets, D12492) at ambient temperature of 22 °C or thermoneutrality (30 °C) for 16 weeks. Food intake and body weight were measured weekly. Fat and lean mass was determined by EchoMRI-100. To measure serum glycerol, FFA and glucose levels of fasted mice, mice were fasted overnight and tail vein blood samples were collected. Free fatty acid levels were measured using a free fatty acid quantitation kit (Sigma, MAK044). Glycerol levels were measured using a free glycerol reagent (Sigma, F6428) or a glycerol cell-based kit (Cayman Chemical, 10011725). To measure serum glycerol, FFA and glucose levels of cold-challenged mice, mice were housed at 4°C overnight. Blood samples were collected from tail veins. To test the effect of OXT on lipolysis *in vivo*, OXT (3mg/kg body weight) and vehicle were injected intraperitoneally into C57BL/6J male mice. To test the effect of OXT on ISO-induced lipolysis in vivo, vehicle, OXT (3mg/kg body weight), ISO (10mg/kg body weight) and OXT+ISO were injected intraperitoneally into C57BL/6J male mice. To test the effect of Atosiban on ISO-induced lipolysis in vivo, vehicle, ISO (10mg/kg body weight) and ISO+Atosiban (5mg/kg body weight) were injected intraperitoneally into C57BL/6J male mice. Serum samples were collected before and 1 hour after injection. To test the effect of OXTR or OXT deficiency on ISO-induced lipolysis, WT and KO mice were injected with ISO (10mg/kg body weight) and blood samples were collected at multiple time points after ISO injection as indicated in the figures.

### *Ex vivo* lipolysis

To test the effect of OXT deficiency in sympathetic neurons innervating iWAT on lipolysis, mice were injected with ISO and iWAT was dissected 20 minutes after ISO injection. iWAT was minced (approximately 0.5mm in diameter), followed by incubation in serum-free DMEM. Samples of media were collected at the time points indicated in the figure and centrifuged at 3000g for 15 min prior to glycerol measurement. To test the effect of OXT on lipolysis and ISO- induced lipolysis, human adipose tissue was minced (approximately 0.5mm in diameter), evenly distributed and cultured in 12-well plates with serum-free DMEM. Vehicle, OXT (Sigma, O3251; 10uM), ISO (10uM) and OXT+ISO with or without Trametinib (Selleck Chemicals, S2673; 5nM) were added to the media. Media samples were collected 4 hours later and centrifuged at 12,000g for 15 minutes prior to glycerol measurement.

### Lipolysis and signal transduction studies in vitro

Primary adipocytes were switched to serum-free DMEM and treated with vehicle, OXT (Sigma, O3251; 10uM), ISO (10uM) or OXT+ISO with or without Erk1/2 signaling inhibitors for 3 hours (for glycerol measurement) or 15 minutes (for signaling studies). To assess dose responsiveness, primary adipocytes were treated with the concentrations of OXT indicated in the figure for 15 minutes. To assess the time course, primary adipocytes were treated with 10uM OXT for the time periods indicated in the figure. To assess the effect of OXT on ISO-induced lipolysis, primary adipocytes were treated with the indicated concentrations of ISO with or without OXT for 3 hours. For measurement of cAMP in primary adipocytes, adipocytes were treated with vehicle, OXT (10uM), ISO (10uM) and OXT+ISO with or without Erk1/2 signaling inhibitors for 15 minutes. cAMP levels were measured from cell lysates using the cAMP Parameter Assay Kit (R&D systems, KGE002B).

### Cold tolerance test

Mice were acclimatized at thermoneutrality (30°C) for four weeks and then shifted to 4°C. Core body temperature was measured using a rectal probe (Yellow Spring Instruments).

### RNA isolation and quantitative PCR

Total RNA from tissues or cells was extracted using Direct-zol RNA MiniPrep kit (ZYMO Research). cDNA was obtained using High-Capacity cDNA Reverse Transcription Kit (Thermo Fisher). RNA levels were measured with the ABI PRISM 7500 (Applied Biosystems). Statistical analysis was performed using ddCt method with 36B4 primers as control. The following primers were used for qPCR: *Oxtr*-forward GCACGGGTCAGTAGTGTCAA, *Oxtr*-reverse AAGCTTCTTTGGGCGCATTG; *Fabp4*-forward TGAAATCACCGCAGACGACA, *Fabp4*- reverse CTCTTGTGGAAGTCACGCCT; *Pparg*-forward TCGCTGATGCACTGCCTATG, *Pparg*-reverse GAGAGGTCCACAGAGCTGATT; *Adipoq*-forward TTGTTCCTCTTAATCCTGCCCA, *Adipoq*-reverse CCAACCTGCACAAGTTCCCTT; *Lipe*- forward GCAGGTGGGAATCTCTGCAT, *Lipe*-reverse GAGGACTGCAGGGTGGTAAC; *Pnpla2*-forward GCATCTCCCTGACTCGTGTT, *Pnpla2*-reverse AATGAGGCCACAGTACACCG; *Oxt*-forward GTGCTGGACCTGGATATGCG, *Oxt*-reverse GGCGAAGGCAGGTAGTTCTC; *36B4-*forward CAGCAGGTGTTTGACAACGG, *36B4-* reverse GATGATGGAGTGTGGCACCG.

### Western blotting

Tissues and cells were homogenized in RIPA buffer containing protease and phosphatase inhibitors (Thermo Fisher) in a Qiagen TissueLyser II (85300). Lysates were then separated by SDS-PAGE, transferred to polyvinylidene fluoride (PVDF) membranes and incubated with primary and secondary antibodies. Antibodies used: mouse monoclonal anti-OXTR (sc-515809), Santa Cruz; anti-HSP90 (#4874), anti-phospho-Erk1/2(#9101), anti-Erk1/2(#4695), anti-Phospho-HSL (Ser660) (#4126), anti-phospho-PKA Substrate (#9624), anti-HSL (#4107), anti-ATGL (#2138), anti-phospho-HSL (Ser565) (#4137), all from Cell Signaling Technology; anti-phospho-Perilipin 1 (Ser522) (Vala Sciences 4856); anti-Perilipin1 (Thermo Fisher PA1-1051); Anti-UCP1 (Abcam ab10983).

### Dissection of relevant sympathetic ganglia

Oxt-Cre::H2B-TRAP mice, Oxt-Cre::Ai9 mice and Oxt-Cre mice injected with AAV2/retro-syn-Flex-mCherry in eWAT or iWAT were deeply anesthetized with an overdose of isoflurane followed by transcardial perfusion with PBS. Thoracic and subdiaphragmatic organs including lung, heart, liver and the gastrointestinal tract were removed to identify paravertebral and prevertebral ganglia and postganglionic neurons. Ganglia with visible fluorescence were dissected using a fluorescent stereomicroscope. Surrounding tissues were removed from sympathetic ganglia prior to further analysis.

### Immunostaining of sympathetic ganglia

Sympathetic ganglia collected from mice were fixed in 10% formalin overnight at 4°C, washed 3 times with PBS and blocked with 5% donkey serum (in 0.4% TBST) for 30 minutes. Sympathetic ganglia were then incubated with primary antibodies for 4 days at 4°C, washed 3 times with PBS and incubated with secondary antibodies for 1.5 hours at room temperature. After washing 4 times with PBS, sympathetic ganglia were mounted with antifade mountant (Invitrogen, P36931). Primary antibodies were: anti-Tyrosine Hydroxylase (Sigma, AB9702), anti-GFP (Abcam, ab290), anti-mCherry (Thermo Fisher Scientific, M11217) and anti-RFP (Rockland, 600-401-379). Secondary antibodies were: Goat 555-conjugated Alexa secondary antibodies against rat (Thermo Fisher Scientific, A-21434), Goat 647-conjugated Alexa secondary antibodies against chicken (Invitrogen, A21449) and Donkey 555-conjugated Alexa secondary antibodies against rabbit (Thermo Scientific, A-31572).

### Immunostaining of primary adipocytes

Primary adipocytes were fixed for 15 min with 4% PFA in PBS, followed by permeabilization for 10 min with 0.1% Triton X-100 in PBS. Adipocytes were then washed with PBS, blocked with 4% donkey serum in PBS for 1 h at RT and incubated with the primary antibodies diluted in blocking solution. Adipocytes were then washed with PBS (3X10 min) and incubated for 1 h at RT with secondary antibodies. Adipocytes were washed with PBS (3X10 min), stained with BODIPY (Thermo Fisher Scientific, D3922) to visualize lipid droplets and mounted in ProLong Gold antifade reagent with DAPI (Invitrogen, P36931). Primary antibodies used were: HSL Antibody (Cell Signaling Technology, 4107) and Perilipin 1 polyclonal antibody (Thermo Fisher Scientific, PA1-1051). The secondary antibodies were Donkey 555-conjugated Alexa secondary antibodies against rabbit (Thermo Scientific, A-31572).

### Retrograde AAV injection

AAV2/retro-hSyn-Cre and AAV2/retro-hSyn-mCherry were gifts from Dr. Bradford Lowell (BIDMC). AAV2/retro-syn-Flex-mCherry was purchased from the Boston Children’s Hospital Viral Core. 8-12 week-old male mice were used in these experiments. For delivery into iWAT, male mice were anesthetized with isoflurane and hair was removed from the inguinal area. A longitudinal incision was made to the skin to expose inguinal adipose tissue. 8 ul AAV was injected into 4 sites of each fat pad using a 10ul Hamilton syringe to distribute the virus to as much of the tissue as possible. The skin was closed with 9mm autoclips, which were removed 2 weeks after injection. In the experiment to knockout *Oxt* specifically in sympathetic neurons innervating iWAT, AAV2/retro-hSyn-mCherry was used as control. Experiments were performed 3 weeks after injection. For delivery into eWAT, laparotomy was performed to expose the eWAT. 4 ul AAV was injected into 2 sites of each fat pad using a 10ul Hamilton syringe (one injection site close to the testicle and the other one in the middle of the fat pad). The abdomen was closed with a two-layer approach.

### Stereotaxic surgery and viral injections

Male mice were anaesthetized with a ketamine (100 mg/kg) and xylazine (10 mg/kg) cocktail diluted in 0.9% saline and mounted into a stereotaxic apparatus (David Kopf model 940). An incision was made to expose the skull and a small craniotomy (coordinate AP/DV/ML: -5.8/-5/0 mm) was made through the skull for virus injection. AAV (50 nL) was injected slowly (20 nL/min) into the medullary raphe via a glass pipette (20–40 um diameter tip) by an air pressure system (Clippard EV 24VDC). The pipette was removed 1 min after each injection and the incision was secured using Vetbond tissue adhesive (3M). Subcutaneous injection of sustained release meloxicam (4 mg/kg) was provided as postoperative care. The AAV2/8-hSyn-DIO-hM3Dq-mCherry (Addgene plasmid 44361; donating investigator, Dr. Bryan Roth) was packaged at BIDMC. The control virus AAV2/5-hSyn-DIOmCherry was ordered from Penn Vector Core (Addgene plasmid 50459). Animals were allowed to recover from stereotaxic surgery for a minimum of 21 days before initiation of any experiments. Accuracy of AAV injections was confirmed via post-hoc histological analysis of mCherry fluorescent protein reporters. All subjects determined to be surgical ‘‘misses’’ based on little or absent reporter expression were removed from further analysis.

### Chemogenetic activation of raphe pallidus

To activate raphe neurons, Saline- or CNO (Sigma, C0832)-loaded minipumps (Braintree scientific, AP-1007D) were implanted subcutaneously to deliver saline or CNO into mice at a constant rate of 0.5ug/hr. Three days later, blood, iWAT, and eWAT samples were collected for analysis.

### Optogenetic activation of sympathetic nerves in eWAT explants

eWAT was dissected from 12-week old male Oxt-Cre::Ai32 mice, minced into small pieces (approximately 0.5mm in diameter), evenly distributed and cultured in 12-well plates with DMEM. The experimental group was exposed to blue light (470nm) for 3 hours, while the control group was exposed to natural light at the same time. Media were collected and oxytocin levels were determined by ELISA.

### Flow cytometry

SVF from adipose tissue of Oxt-Cre::Ai9 mice and Ai9 mice was isolated as described above. SVF cells were suspended in PBS containing 1% BSA and filtered through 20uM cell strainers. Flow cytometry was performed using a BD FACS Aria II, gating on FSC, SSC, and DsRED fluorescence in the BIDMC FACS core facility.

### Adipo-clear

Adipo-Clear was performed as described^28,38^. Briefly, mice were heavily anesthetized with an overdose of isoflurane and intracardiac perfusion and fixation was performed with PBS followed by 4% PFA. All harvested samples were post-fixed in 4% PFA at 4°C overnight. Fixed samples were washed in PBS for 1 hr three times, then washed in 20%, 40%, 60%, 80% methanol in H_2_O/0.1% Triton X-100/0.3M glycine (B1N buffer, pH 7), and 100% methanol for 30 min each. Sample were then delipidated with 100% dichloromethane (DCM; Sigma-Aldrich) for 30 min three times. After delipidation, samples were washed in 100% methanol for 30 min twice, then in 80%, 60%, 40%, 20% methanol in B1N buffer for 30 min at each step. All procedures above were carried out at 4°C with shaking. Samples were then washed in B1N for 30 min twice followed by PBS/0.1% Triton X-100/0.05% Tween 20/2 ug/ml heparin (PTwH buffer) for 1hr twice before further staining procedures. For immunolabeling, samples were incubated in primary antibody dilutions in PTxwH for 4 days. After primary antibody incubation, samples were washed in PTxwH for 5 min, 10 min, 15 min, 30 min, 1 hr, 2 hr, 4 hr, and overnight, and then incubated in secondary antibody dilutions in PTxwH for 4 days. Samples were finally washed in PTxwH for 5 min, 10 min, 15 min, 30 min, 1 hr, 2 hr, 4 hr, and overnight. In this study, GFP (1:500, Aves Labs, GFP-1020) and TH (1:200, Millipore, AB152) were used. Secondary antibodies conjugated with Alexa-568 (Invitrogen, A10042) and Alexa-647 (Jackson ImmunoResearch, 703-605-155) were used (1:200). Samples were dehydrated in 25%, 50%, 75%, 100%, 100% methanol/H_2_O series for 30 min at each step at RT. Following dehydration, samples were washed with 100% DCM for 30 min twice, followed by an overnight clearing step in dibenzyl ether (DBE; Sigma-Aldrich). Samples were stored at RT in the dark until imaging. Samples were imaged on a light-sheet microscope (Ultramicroscope II, LaVision Biotec) equipped with a 4X objective and an sCMOs camera (Andor Neo). Images were acquired with the ImspectorPro software (LaVision BioTec). Samples were placed in an imaging reservoir filled with DBE and illuminated from the side by the laser light sheet. The samples were scanned with the 488, 561, and 640nm laser channels and with a step-size of 2.5 µ m. Images were generated using Imaris x64 software (Bitplane). 3D reconstruction was performed using the ‘‘volume rendering’’ function. Optical slices were obtained using the ‘‘orthoslicer’’ tool. Optical sections were generated using the ‘‘snapshot’’ tool.

### Brain tissue preparation

Animals were terminally anesthetized with 7% chloral hydrate diluted in saline (350 mg/kg) and transcardially perfused with PBS (10 mL) then 10% neutral-buffered formalin (∼15 mL). Brains were removed, stored in the same fixative overnight, transferred into 20% sucrose at 4°C overnight, and sectioned into 30 μm sections on a freezing microtome (Leica Biosystems) coronally into three equal series and stored at -20°C in antifreeze solution (25% ethylene glycol, 25% glycerol in PBS).

### Immunohistochemistry

Brain sections were washed with PBS and then placed in blocking solution (PBS containing 0.1% TritonX-100, 4% normal goat serum, Jackson Immunoresearch) for 1 hr at room temperature. For TH and Oxytocin colocalization, brain sections containing PVH and SON were incubated with sheep anti-Tyrosine Hydroxylase (AB152, Millipore, 1:2000) and rabbit anti-Oxytocin (#20068 Immunostar, 1:1000) in blocking solution for 48 hr at 4°C; for cFos activation in DREADDs experiments, brain sections containing raphe were incubated with rabbit anti-cFos (Abcam #ab214672, 1:1000) and rat anti-mCherry (Invitrogen M11217, 1:5000). Sections were then washed 4X in PBS, then incubated for 2 hours at room temperature in Alexa Fluor fluorescent secondary antibody (Life Technologies; 1:500) prepared in blocking solution. Finally, sections were washed 4X in PBS, mounted on gelatin-coated slides, and coverslipped (VWR International 48393 251) with ProLong™ Gold Antifade Mountant containing DAPI (P10144, Molecular Probe). Fluorescent images were captured using an Olympus VS120 slide-scanning microscope.

### Statistical analysis

Wild-type mice were randomly assigned to treatment groups. No animal experiment was blinded. All data in this study were normally distributed, and no data points were excluded from analysis. Results are shown as mean ± s.e.m. Statistical analysis was performed using Prism (GraphPad). Comparisons between the two groups were analysed using the 2-tailed paired Student’s t test or the Student’s t test, as indicated in figure legends. Two-way ANOVA with Tukey post hoc analysis was used for multiple group comparisons. P<0.05 was considered statistically significant. Asterisks denote corresponding statistical significance *p < 0.05. The definition of ‘‘n’’ is indicated in the figure legends.

## ACKNOWLEDGEMENTS

This work was supported by NIH grants R01 DK126789 to EDR, P30 DK056341 to SK, and R01 DK120649 to PC. We thank Christina Usher for artistic support. We thank Heike Muenzberg-Gruening for helpful discussions, and the study subjects for their participation.

## AUTHOR CONTRIBUTIONS

EL and EDR conceived of the project. EL, LW, DW, GIS, and JC performed experiments. SK and GIS provided human samples. EL and EDR wrote the manuscript; all authors reviewed the manuscript prior to submission.

## ADDITIONAL INFORMATION

Supplementary information is available for this paper.

Correspondence and requests for materials should be addressed to EDR.

---

**Extended Data Fig. 1.**
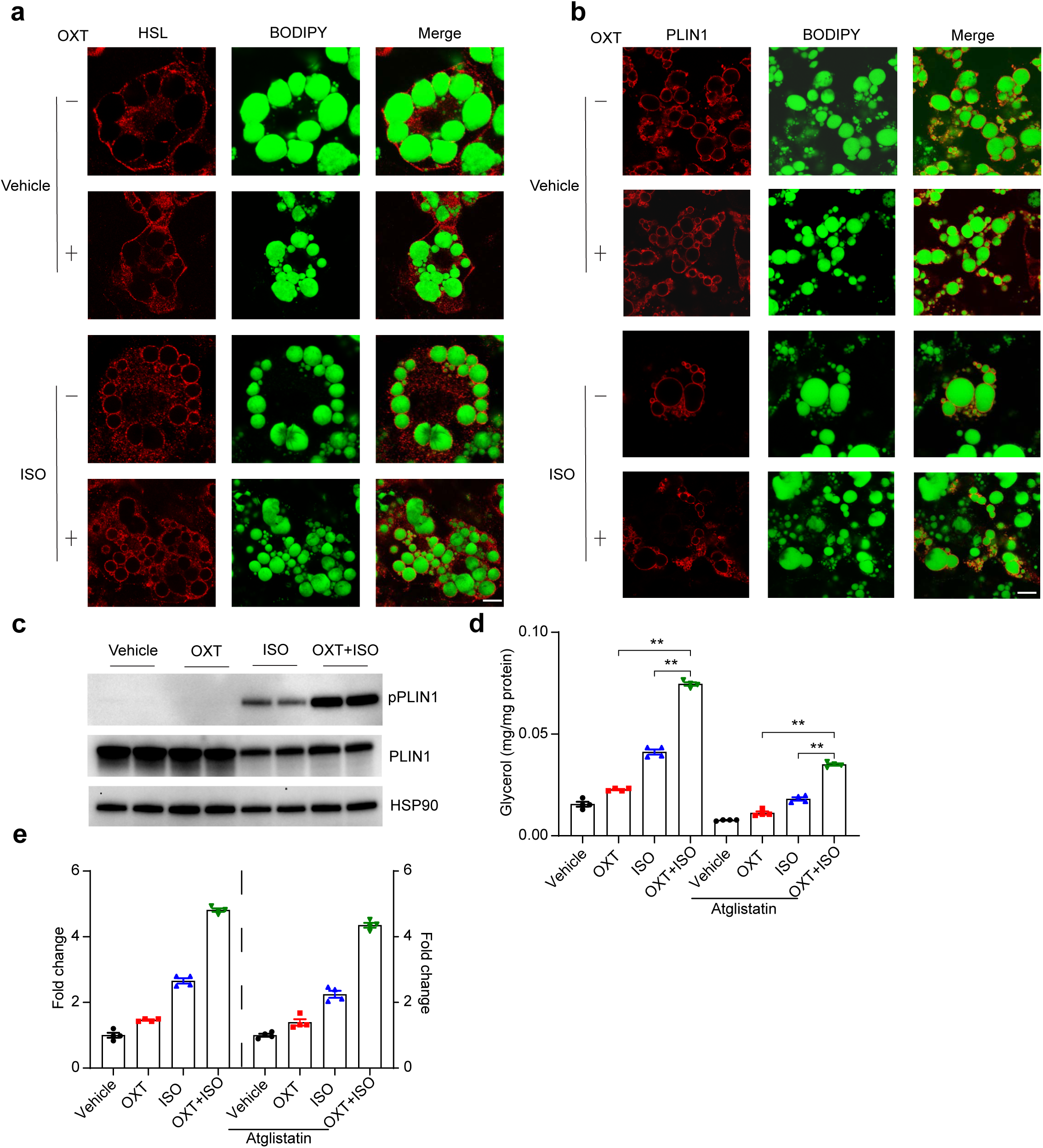
OXTR expression in adipocytes. **a.** *Oxtr* mRNA expression in mouse eWAT, iWAT and BAT; n=4. **b.** OXTR protein expression in mouse eWAT, iWAT and BAT. Representative image from two western blots. **c.** *Oxtr* mRNA expression in the SVF and adipocytes of mouse iWAT and eWAT; n=4. **d.** OXTR protein expression in the SVF and adipocytes of mouse iWAT and eWAT. Representative image from two western blots. **e.** *Oxtr* mRNA expression in SVF and adipocytes differentiated from SVF ex vivo; n=4. **f.** OXTR protein levels in SVF and adipocytes differentiated from SVF ex vivo. Representative image from two western blots. **g.** *Oxtr* mRNA expression by qPCR in isolated adipocytes of iWAT and eWAT, harvested from mice housed at 30°C or 4°C for 1 week; warm n=5, cold n=6. **h.** OXTR protein expression in isolated adipocytes of iWAT and eWAT, harvested from mice housed at 30°C or 4°C for 1 week. Representative image from two western blots. Data are presented as mean ± s.e.m. Statistical comparisons were made using 2-tailed Student’s t test. *P < 0.05; **P < 0.01.

**Extended Data Fig. 2.**
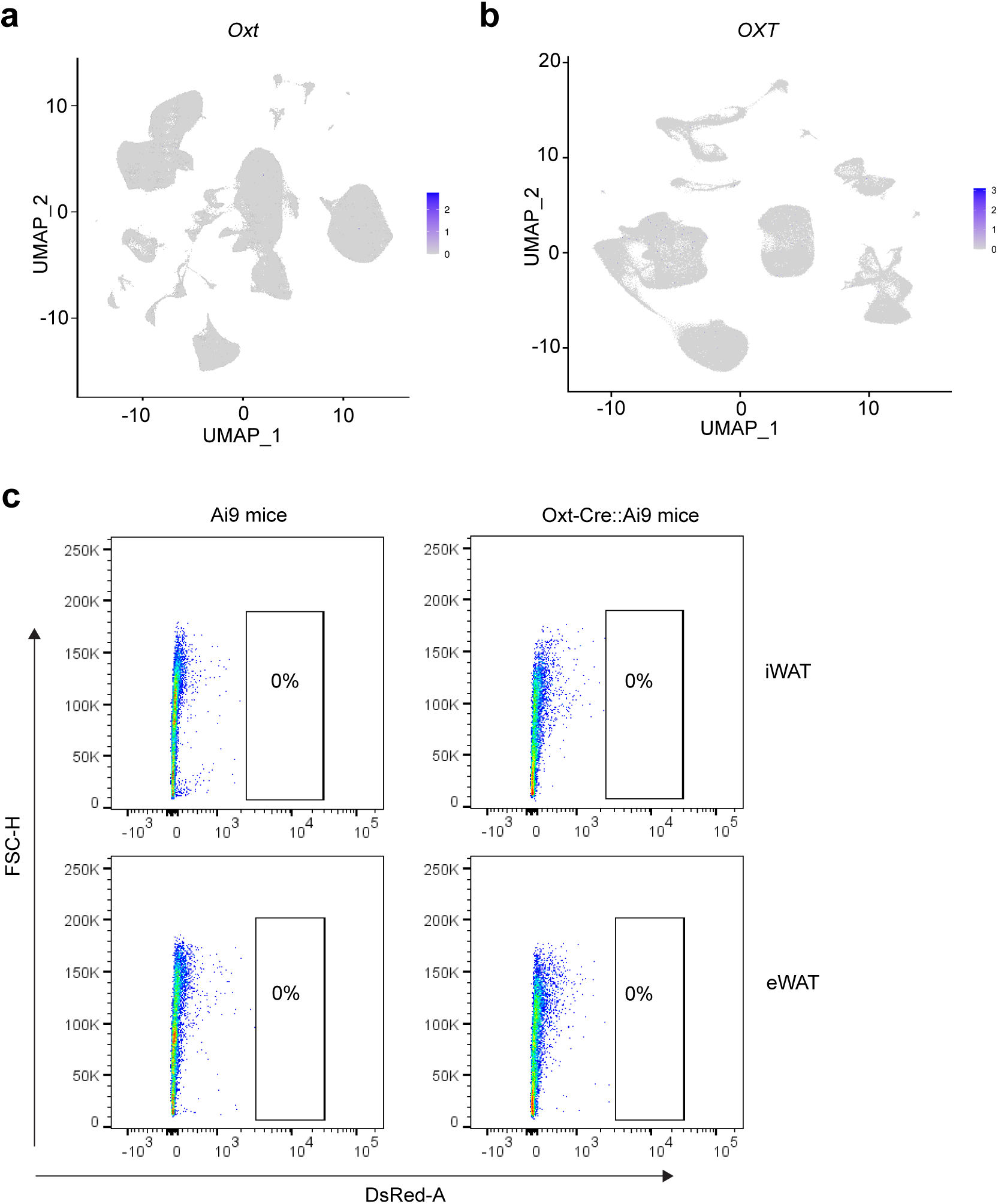
OXT induces lipolysis through ERK signaling. **a.** Glycerol release from cultured adipocytes treated with different doses of OXT for 3 hours; n=3. **b.** cAMP levels in cultured adipocytes treated with vehicle, 1uM or 10 uM OXT for 15 minutes; n=3. **c.** Western blotting of PKA substrate from cultured adipocytes treated with vehicle or 10uM OXT for 15 minutes; n=3. **d.** Glycerol release from cultured adipocytes treated with vehicle or ISO in the presence of DMSO, PKG inhibitor KT5823 or PKA inhibitor H89; n=3. **e.** Glycerol release from cultured adipocytes treated with vehicle or OXT in the presence of DMSO, PKG inhibitor KT5823 or PKA inhibitor H89; n=3. **f.** Dose response for pERK1/2 induced by OXT in cultured adipocytes. Representative image from two western blots. **g.** Time course of OXT-induced (10uM) ERK activation and PKA activity in cultured adipocytes. Representative image from two western blots. **h.** Glycerol release from cultured adipocytes treated with OXT (10uM), OXT + the MEK inhibitor Trametinib (Tra)(5 nM) or OXT + the ERK inhibitor Temuterkib (Tem)(2 uM); n=3. Data are presented as mean ± s.e.m. Statistical comparisons were made using 2-tailed Student’s t test. *P < 0.05; **P < 0.01.

**Extended Data Fig. 3.**
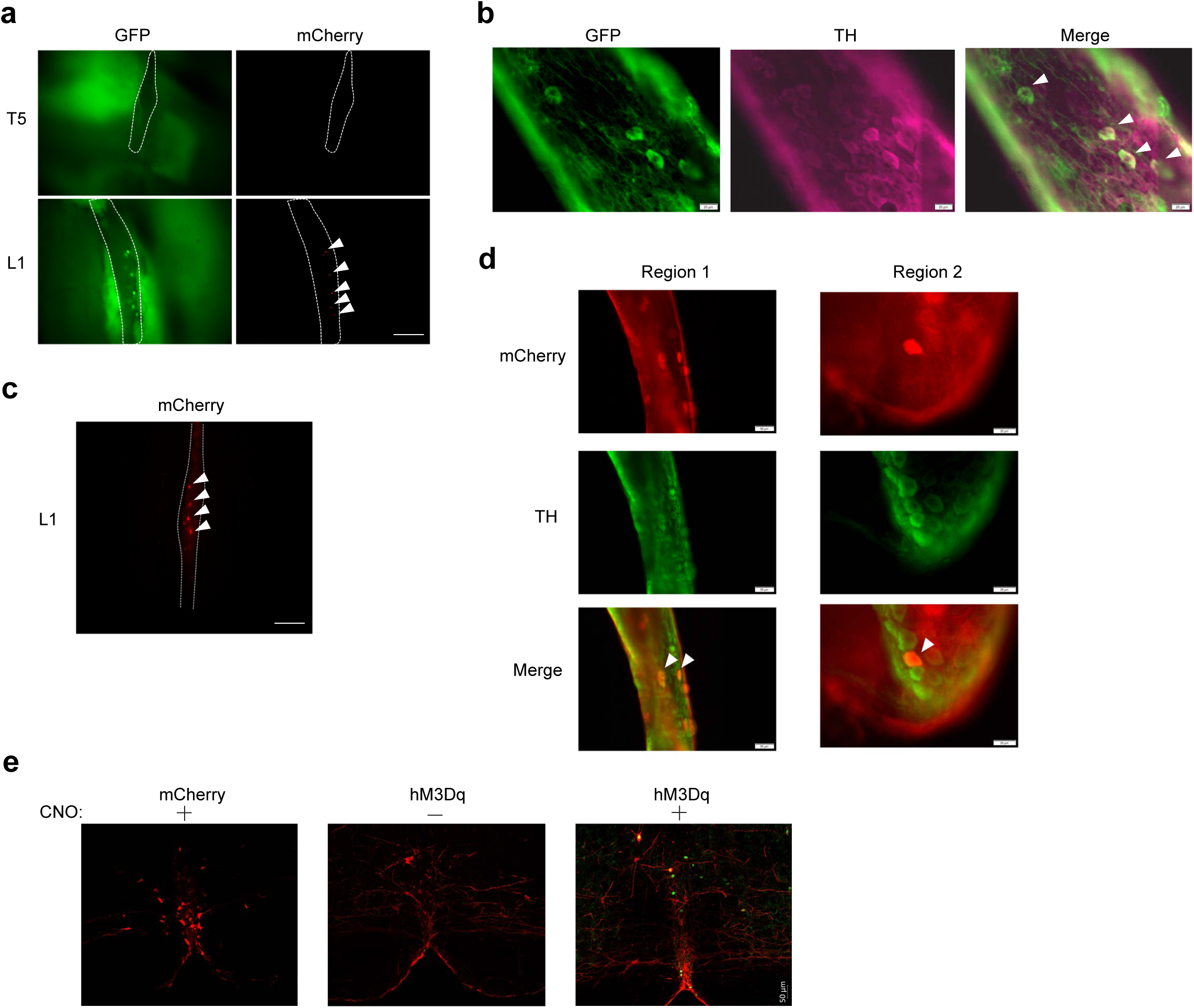
Knockout of OXTR in adipocytes has minimal effect on body weight. **a.** Scheme of the floxed *Oxtr* allele. **b.** *Oxtr* mRNA levels in adipocytes of eWAT and iWAT from *Oxtr*^flox^ and *Oxtr*^ΔAd^ mice housed at 4°C for 1 week; n=5. **c.** Oxtr protein levels in adipocytes of eWAT from *Oxtr*^flox^ and *Oxtr*^ΔAd^ mice housed at 4°C for 1 week; n=3. **d.** Body weight of the *Oxtr*^flox^ and *Oxtr*^ΔAd^ mice on chow and HFD; n=8-15. **e.** Lean and fat mass of the *Oxtr*^flox^ and *Oxtr*^ΔAd^ mice on chow; n=8-10. **f.** Tissue weight of liver, BAT, iWAT, and eWAT of *Oxtr*^flox^ and *Oxtr*^ΔAd^ mice on HFD for 16 weeks; n=7. **g.** H&E staining of liver, BAT, iWAT, and eWAT of *Oxtr*^flox^ and *Oxtr*^ΔAd^ mice on HFD for 16 weeks. Scale bar, 200 um. **h.** Rectal temperature of *Oxtr*^flox^ and *Oxtr*^ΔAd^ mice exposed to 4°C; n=5-6. **i.** Western blotting of UCP-1 in BAT and iWAT of chow-fed *Oxtr*^flox^ and *Oxtr*^ΔAd^ mice housed at 4°C for 1 week; n=4. **j.** mRNA levels of thermogenic genes in cultured adipocytes treated with vehicle or OXT (10uM) for 24 hours; n=3. **k.** Western blotting of UCP-1, HSL and ATGL in cultured adipocytes treated with vehicle or OXT (10uM) for 24 hours. Representative image from two western blots. Data are presented as mean ± s.e.m. Statistical comparisons were made using 2-tailed Student’s t test (b, e, f, j) or 2-way ANOVA (d, h). *P < 0.05; **P < 0.01.

**Extended Data Fig. 4.**
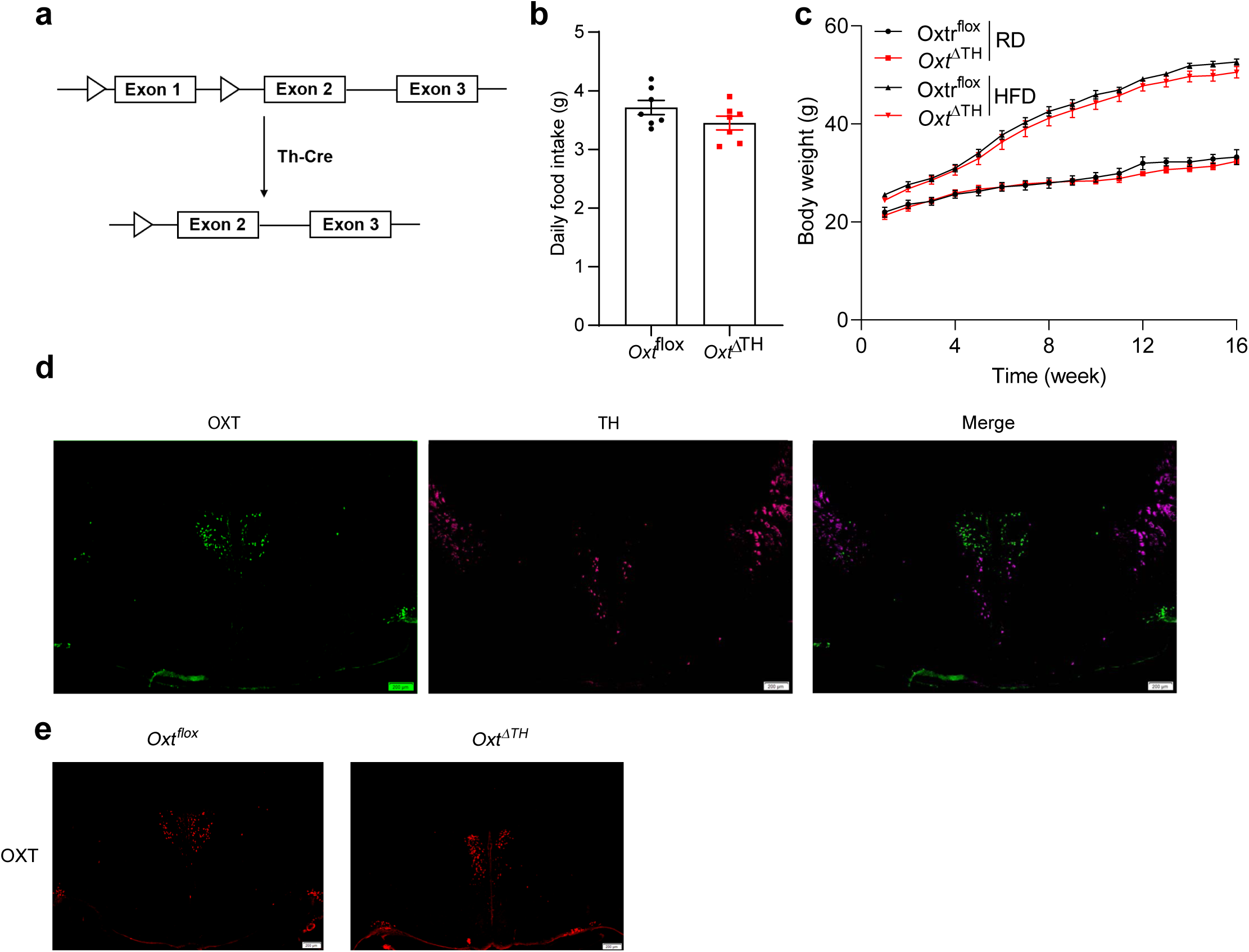
Effect of OXT on ISO-induced lipolysis. **a.** *Lipe* and *Pnpla2* mRNA levels in eWAT and iWAT of chow-fed *Oxtr*^flox^ and *Oxtr*^ΔAd^ mice housed at 4°C for 1 week; n=5-6. **b.** Western blotting of pPLIN1, PLIN1, pHSL, HSl, and ATGL of eWAT and iWAT of *Oxtr*^flox^ and *Oxtr*^ΔAd^ mice housed at 4°C for 1 week; n=5. **c.** Glycerol release from human adipose explants treated with OXT (10uM), isoproterenol (ISO; 10uM), or both in the presence or absence of the MEK inhibitor trametinib (Tra) (5 nM); n=4. **d.** Glycerol release from human adipose explants treated with different doses of ISO with or without OXT (10uM); n=4. **e.** Western blotting of pPLIN1, pHSL, pERK, and PKA activity from the same cells shown in Fig. 1f. Representative image from two western blots. **f.** cAMP levels in the same cells shown in Fig. 1f; n=3. Data are presented as mean ± s.e.m. Statistical comparisons were made using 2-tailed Student’s t test (a, c, f) or 2-way ANOVA (d). *P < 0.05; **P < 0.01.

**Extended Data Fig. 5.**
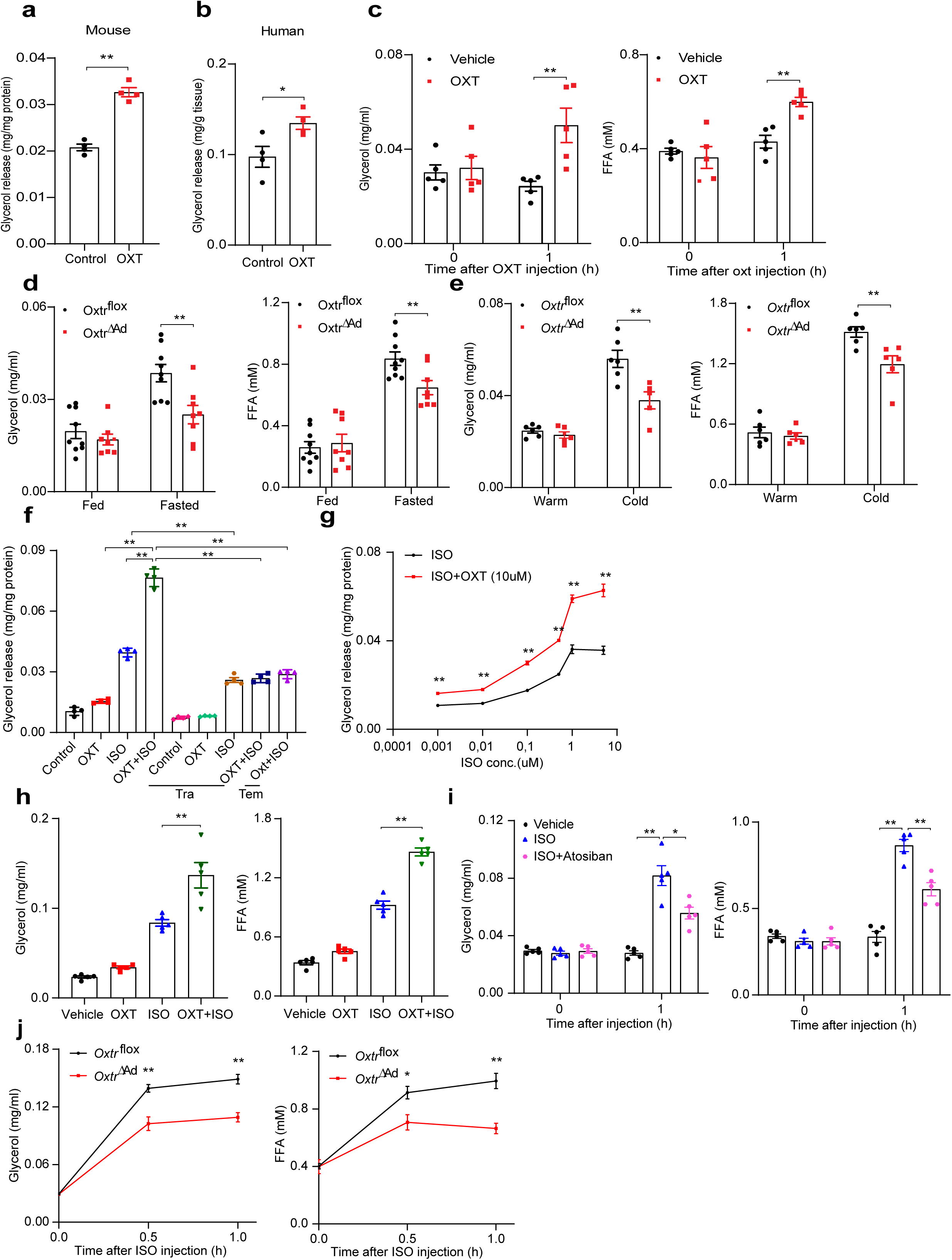
Effect of OXT on HSL, PLIN1 and ATGL. **a.** Representative images showing the distribution of HSL in cultured adipocytes treated with vehicle or OXT (10uM) in the presence or absence of ISO (10uM). Scale bar, 10 um. **b.** Representative images showing the distribution of PLIN1 in lipid droplets of cultured adipocytes treated with vehicle or OXT (10uM) in the presence or absence of ISO (10uM). Scale bar, 20 um. **c.** Western blotting of pPLIN1 and PLIN1 in cultured adipocytes treated with OXT (10uM), ISO (10uM), or both for 3 hours. Representative image from two western blots. **d.** Glycerol release from cultured adipocytes treated with OXT (10uM), ISO (10uM), or both in the presence or absence of the ATGL inhibitor Atglistatin (100uM) for 3 hours; n=4. **e.** Fold change of glycerol release in **d** (normalized to control); n=4. Data are presented as mean ± s.e.m. Statistical comparisons were made using 2-tailed Student’s t test. *P < 0.05; **P < 0.01.

**Extended Data Fig. 6.**
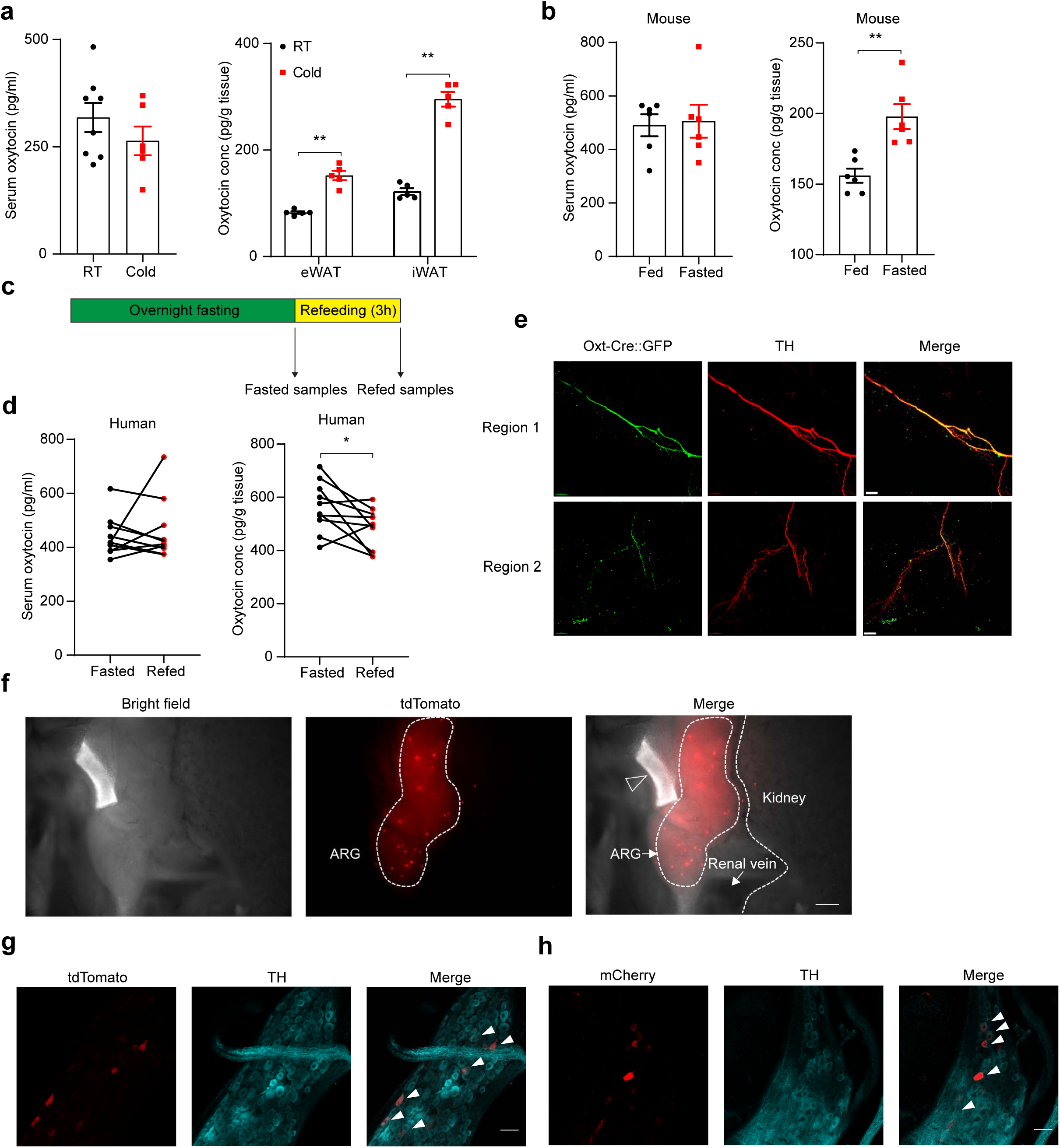
No cells make OXT in adipose tissue. **a.** Single nuclei RNA-seq of mouse adipose tissue (epididymal and inguinal) shows undetectable *Oxt* expression. **b.** Single cell RNA-seq of human white adipose tissue (subcutaneous and visceral) shows undetectable *OXT* expression. **c.** Representative flow cytometric analysis of tdTomato-positive cells in iWAT and eWAT SVF of Oxt-Cre::Ai9 mice.

**Extended Data Fig. 7.**
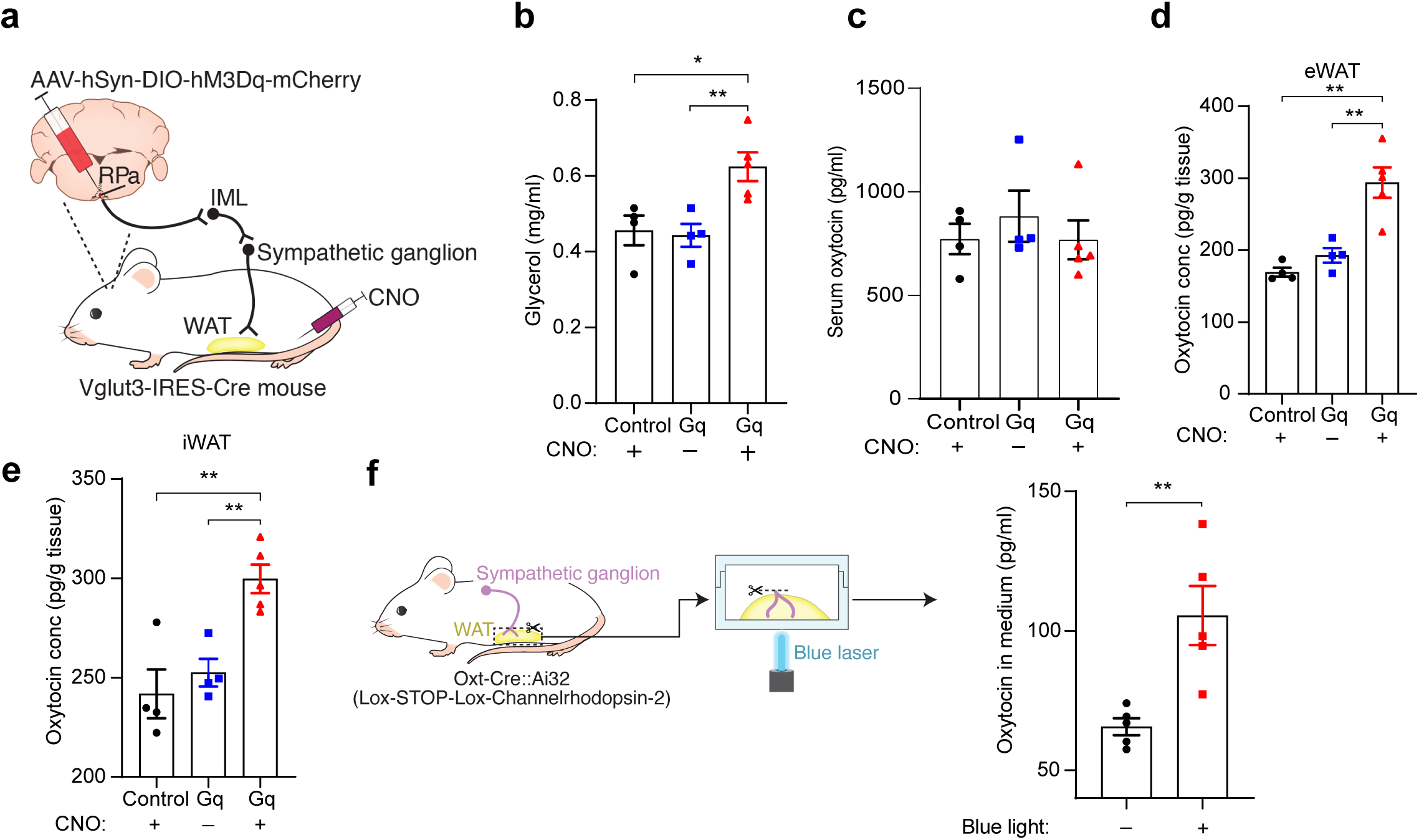
Sympathetic nerves are the source of OXT in fat. **a.** GFP-positive cells and mCherry-positive nuclei in L5 and L1 ganglia of Oxt-Cre::H2B-TRAP mice. Scale bar, 200 um. **b.** Co-immunostaining of GFP and TH in the L1 ganglion of *Oxt*-Cre::H2B-TRAP mice. Scale bar, 20 um. **c.** mCherry-positive cells in the L1 ganglion of Oxt-Cre mice injected with AAV2/retro-syn- Flex-mCherry in iWAT. Scale bar, 200 um. **d.** Co-immunostaining of mCherry and TH in the L1 ganglion of mice in **c**. Scale bars, 50 um (region 1) and 20 um (region 2). **e.** RPa from mice in Fig. 3b, showing mCherry in all mice, but c-Fos activation (green) in mice that received both DREADD and CNO. Scale bar, 50 um.

**Extended Data Fig. 8.**
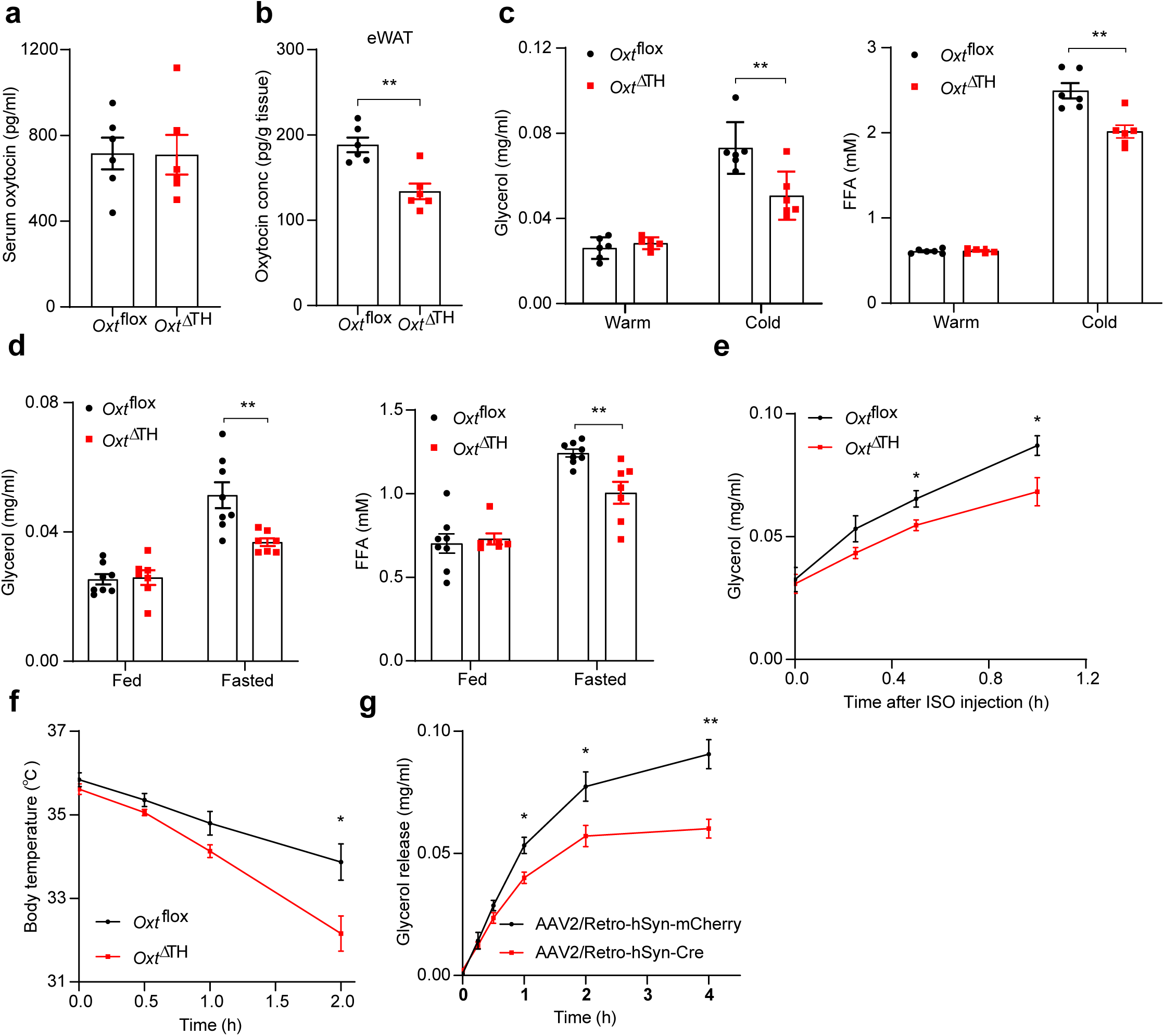
Loss of OXT in TH+ neurons reduces lipolysis. **a.** Scheme of the floxed *Oxt* allele. **b.** Daily food intake of chow-fed *Oxt*^flox^ and *Oxt*^ΔTH^ mice; n=7. **c.** Body weight of *Oxt*^flox^ and *Oxt*^ΔTH^ mice on chow and HFD; n=7-10. **d.** Co-immunostaining of OXT and TH in PVN and SON of WT mice. Scale bar, 200 um. **e.** Immunostaining of OXT in PVN of *Oxt*^flox^ and *Oxt*^ΔTH^ mice. Scale bar, 200 um. Data are presented as mean ± s.e.m. Statistical comparisons were made using 2-tailed Student’s t test (b) or 2-way ANOVA (c). *P < 0.05; **P < 0.01.

## REFERENCES

1. Lawson, E. A. The effects of oxytocin on eating behaviour and metabolism in humans. Nature Reviews Endocrinology 13, 700–709 (2017).

2. Romano, A., Tempesta, B., Micioni Di Bonaventura, M. V. & Gaetani, S. From autism to eating disorders and more: the role of oxytocin in neuropsychiatric disorders. Frontiers in neuroscience 9, 497 (2016).

3. McCormack, S. E., Blevins, J. E. & Lawson, E. A. Metabolic effects of oxytocin. Endocrine reviews 41, 121–145 (2020).

4. Ding, C., Leow, M. S. & Magkos, F. Oxytocin in metabolic homeostasis: implications for obesity and diabetes management. Obesity Reviews 20, 22–40 (2019).

5. Blevins, J. E. et al. Chronic oxytocin administration inhibits food intake, increases energy expenditure, and produces weight loss in fructose-fed obese rhesus monkeys. *American Journal of Physiology-Regulatory*, Integrative and Comparative Physiology 308, R431–R438 (2015).

6. Blevins, J. E. & Baskin, D. G. Translational and therapeutic potential of oxytocin as an anti- obesity strategy: Insights from rodents, nonhuman primates and humans. Physiology & behavior 152, 438–449 (2015).

7. Deblon, N. et al. Mechanisms of the anti-obesity effects of oxytocin in diet-induced obese rats. PloS one 6, e25565 (2011).

8. Burt, R. L., Leake, N. H. & Dannenburg, W. N. Metabolic activity of oxytocin in the puerperium. Nature 198, 293–293 (1963).

9. Sun, L. et al. Oxytocin regulates body composition. Proceedings of the National Academy of Sciences 116, 26808–26815 (2019).

10. Grabner, G. F., Xie, H., Schweiger, M. & Zechner, R. Lipolysis: cellular mechanisms for lipid mobilization from fat stores. Nature Metabolism 3, 1445–1465 (2021).

11. Sengenès, C., Berlan, M., de Glisezinski, I., Lafontan, M. & Galitzky, J. Natriuretic peptides: a new lipolytic pathway in human adipocytes. The FASEB Journal 14, 1345–1351 (2000).

12. Jurek, B. & Neumann, I. D. The oxytocin receptor: from intracellular signaling to behavior. Physiological reviews 98, 1805–1908 (2018).

13. Iovino, M. et al. Oxytocin Signaling Pathway: From Cell Biology to Clinical Implications. Endocrine, Metabolic & Immune Disorders-Drug Targets (Formerly Current Drug Targets- Immune, Endocrine & Metabolic Disorders) 21, 91–110 (2021).

14. El-Merahbi, R. et al. The adrenergic-induced ERK3 pathway drives lipolysis and suppresses energy dissipation. Genes & development 34, 495–510 (2020).

15. Zong, J. et al. Bromodomain-containing protein 2 promotes lipolysis via ERK/HSL signalling pathway in white adipose tissue of mice. General and comparative endocrinology 281, 105–116 (2019).

16. Lee, H.-J., Caldwell, H. K., Macbeth, A. H., Tolu, S. G. & Young 3rd, W. S. A conditional knockout mouse line of the oxytocin receptor. Endocrinology 149, 3256–3263 (2008).

17. Eguchi, J. et al. Transcriptional control of adipose lipid handling by IRF4. Cell metabolism 13, 249–259 (2011).

18. Greenberg, A. S. et al. Stimulation of lipolysis and hormone-sensitive lipase via the extracellular signal-regulated kinase pathway. Journal of Biological Chemistry 276, 45456–45461 (2001).

19. Su, C.-L. et al. Mutational analysis of the hormone-sensitive lipase translocation reaction in adipocytes. Journal of Biological Chemistry 278, 43615–43619 (2003).

20. Greenberg, A. S. et al. Perilipin, a major hormonally regulated adipocyte-specific phosphoprotein associated with the periphery of lipid storage droplets. Journal of Biological Chemistry 266, 11341–11346 (1991).

21. Miyoshi, H. et al. Perilipin promotes hormone-sensitive lipase-mediated adipocyte lipolysis via phosphorylation-dependent and-independent mechanisms. Journal of Biological Chemistry 281, 15837–15844 (2006).

22. Tansey, J. et al. Perilipin ablation results in a lean mouse with aberrant adipocyte lipolysis, enhanced leptin production, and resistance to diet-induced obesity. Proceedings of the National Academy of Sciences 98, 6494–6499 (2001).

23. Emont, M. P. et al. A single-cell atlas of human and mouse white adipose tissue. Nature, 1–8 (2022).

24. Brito, N. A., Brito, M. N. & Bartness, T. J. Differential sympathetic drive to adipose tissues after food deprivation, cold exposure or glucoprivation. American Journal of Physiology-Regulatory, Integrative and Comparative Physiology 294, R1445–R1452 (2008).

25. Ohlsson, B., Truedsson, M., Djerf, P. & Sundler, F. Oxytocin is expressed throughout the human gastrointestinal tract. Regulatory peptides 135, 7–11 (2006).

26. Dayanithi, G. et al. Vasopressin and oxytocin in sensory neurones: expression, exocytotic release and regulation by lactation. Scientific reports 8, 1–12 (2018).

27. Roh, H. C. et al. Simultaneous transcriptional and epigenomic profiling from specific cell types within heterogeneous tissues in vivo. Cell reports 18, 1048–1061 (2017).

28. Chi, J. et al. Three-dimensional adipose tissue imaging reveals regional variation in beige fat biogenesis and PRDM16-dependent sympathetic neurite density. Cell metabolism 27, 226–236. e223 (2018).

29. Huesing, C. et al. Sympathetic innervation of inguinal white adipose tissue in the mouse. Journal of Comparative Neurology 529, 1465–1485 (2021).

30. Cardoso, F. et al. Neuro-mesenchymal units control ILC2 and obesity via a brain–adipose circuit. Nature 597, 410–414 (2021).

31. François, M., Qualls-Creekmore, E., Berthoud, H.-R., Münzberg, H. & Yu, S. Genetics-based manipulation of adipose tissue sympathetic innervation. Physiology & behavior 190, 21–27 (2018).

32. Klein, S., Sakurai, Y., Romijn, J. A. & Carroll, R. M. Progressive alterations in lipid and glucose metabolism during short-term fasting in young adult men. American Journal of Physiology- Endocrinology And Metabolism 265, E801–E806 (1993).

33. Horowitz, J. F. & Klein, S. Lipid metabolism during endurance exercise. The American journal of clinical nutrition 72, 558S–563S (2000).

34. Petersen, M. C., Vatner, D. F. & Shulman, G. I. Regulation of hepatic glucose metabolism in health and disease. Nature reviews endocrinology 13, 572–587 (2017).

35. Haemmerle, G. et al. Defective lipolysis and altered energy metabolism in mice lacking adipose triglyceride lipase. Science 312, 734–737 (2006).

36. Wueest, S. et al. Mesenteric fat lipolysis mediates obesity-associated hepatic steatosis and insulin resistance. Diabetes 65, 140–148 (2016).

37. Niu, J., Tong, J. & Blevins, J. E. Oxytocin as an Anti-obesity Treatment. Frontiers in Neuroscience, 1325 (2021).

38. Chi, J., Crane, A., Wu, Z. & Cohen, P. Adipo-clear: a tissue clearing method for three- dimensional imaging of adipose tissue. JoVE (Journal of Visualized Experiments), e58271 (2018).

